# Meta-analysis and multi-omics to elucidate pathogenic mechanisms of age-related knee osteoarthritis

**DOI:** 10.1101/2021.05.06.442993

**Authors:** Hirotaka Iijima, Gabrielle Gilmer, Kai Wang, Sruthi Sivakumar, Christopher Evans, Yusuke Matsui, Fabrisia Ambrosio

## Abstract

Increased mechanistic insight into the pathogenesis of knee osteoarthritis (KOA) is needed to develop efficacious disease-modifying treatments. Though age-related pathogenic mechanisms are most relevant to the great majority of KOA seen clinically, the bulk of our mechanistic understanding of KOA has been derived using surgically induced post-traumatic OA (PTOA) models. Here, we took an integrated approach of meta-analysis and multi-omics to elucidate pathogenic mechanisms of age-related KOA in murine model. Protein-level data together with transcriptomic profiling revealed inflammation, autophagy, and cellular senescence as primary hallmarks of age-related KOA. Importantly, the molecular profiles of aged cartilage were unique from those in PTOA, with only 1% overlap between the two. At the nexus of aging hallmarks, Advanced Glycation End-Product (AGE)/Receptor for AGE emerged as intrinsically linked to age-related KOA. This pathway was further validated by mass spectrometry. Collectively, these findings implicate dysregulation of AGE-RAGE signaling as a key driver of age-related KOA.

## INTRODUCTION

“ *What doesn’t kill you, makes your stronger*.” In 1888, Friedrich Nietzsche made this proclamation; a proclamation that has echoed long beyond his years, perhaps because it resonates with our inherent desire to not only survive but to thrive. Over the last century, medical research has championed increasing the lifespan of our population, with encouraging success. However, a new set of epidemic diseases have made clear the all-too-common disconnect between surviving and thriving. One of the most debilitating, yet non-lethal, diseases is knee osteoarthritis (KOA), which represented the 12th leading cause of disability worldwide in 2016(1). The number of years lived with KOA-induced disability increased by 31.5% from 2006 to 2016(1), suggesting the impact of KOA on society is rapidly expanding. Yet, progress in developing effective interventions for KOA has been slow, with current therapeutic strategies primarily focusing on symptom management, such as pain reduction and compensatory approaches to improve physical function(2). The development of effective and sustainable disease-modifying treatment is, therefore, a critical challenge to reduce the economic burden of KOA and extend the healthspan of our aging population.

Several clinical trials targeting molecules implicated in the pathogenesis of KOA have been tested, though none have proven effective. For example, inducible nitric oxide synthase (iNOS) has been shown in a post-traumatic OA (PTOA) animal model to be a key driver of cartilage damage(3). Unfortunately, clinical translation of drugs targeting this molecule was unsuccessful, as evidenced by a lack of structural improvement in older subjects with KOA(4). Likewise, inhibitors of a-disintegrin and metalloproteinase with thrombospondin motifs (ADAMTS) have also been investigated due to the reported role of these enzymes in the degradation of proteoglycans in PTOA animal model(5). However, most phase I and II clinical trials involving ADAMTS have not reported findings and/or have ceased further development(6). The lack of success in these trials is attributed, at least in part, to our inadequate understanding of the cellular and molecular mechanisms driving KOA.

*What has impeded progress towards mechanistic understanding of KOA pathogenesis?* Due to challenges obtaining longitudinal human OA cartilage samples and difficulties securing valid and reliable control samples, animal models are a staple of KOA mechanistic studies. In these efforts, animal models where OA is induced surgically or mechanically (i.e., PTOA) are most commonly investigated owing to their convenience and reproducibility. However, the translatability of findings from PTOA animal models to age-related OA in humans is unclear. For instance, unlike in PTOA models, most patients with OA do not present with a clear inciting event. Instead, aging is the single greatest predictive factor for KOA. PTOA, on the other hand, only accounts for 12% of the total KOA burden(7). The study of natural-aging KOA models is therefore imperative for enhanced mechanistic insight into the disease process in the majority of humans.

Meta-analysis of the literature allows for both resolution of inconclusive results and identification of novel candidate pathways promotes a more comprehensive understanding of the disease under study. The strength of meta-analysis is further enhanced when identified candidate pathways are validated by multi-scale omics data. In this study, we took the novel approach of integrated meta-analysis together with transcriptomic and proteomic bioinformatics with the goal of identifying common molecular denominators of age-related KOA. Specifically, we first performed a systematic literature review to identify all relevant age-related KOA studies in murine models and compiled findings via meta-analysis. We then integrated protein data from included studies in order to determine novel pathways associated with KOA and validated the identified pathways by mass spectrometry-based proteomic analysis. Next, we cross-checked the identified pathways with RNA-seq data and merged transcriptomic and protein data to further understand the levels of regulation within KOA. Lastly, we evaluated the methodological rigor and reproducibility of the included studies following the previously established reporting guideline, Animal Research: Reporting of In Vivo Experiments (ARRIVE)(8).

## RESULTS

A systematic search identified a total of 1,222 articles from electronic databases relating to KOA and aging, of which 40 met inclusion criteria for this systematic review (**Figure 1, Table S1**). The rationale for excluding studies during the full-text screening process is provided in **Table S2**. The most common reason for exclusion was the lack of an “ aged” comparison group over 18 months old. **Table S3** summarizes the characteristics of the included studies.The most commonly used murine strains were from a C57BL/6 background, and the median age of young, middle-aged, and aged mice was 5 months (interquartile (IQ) range: 3-6), 12 months (IQ range: 12-12), and 22 months (IQ range: 18-24), respectively. Remarkably, of the 29 studies that reported the sex of the mice, only 6 studies included both male and female mice. The remaining 23 studies used either only male (22 studies) or female (1 study) mice. This is a critical gap in the literature given the predominance of OA in women after menopause(9). Outcome measurements in the included studies primarily examined the tibiofemoral joint.

**Figure 1.**
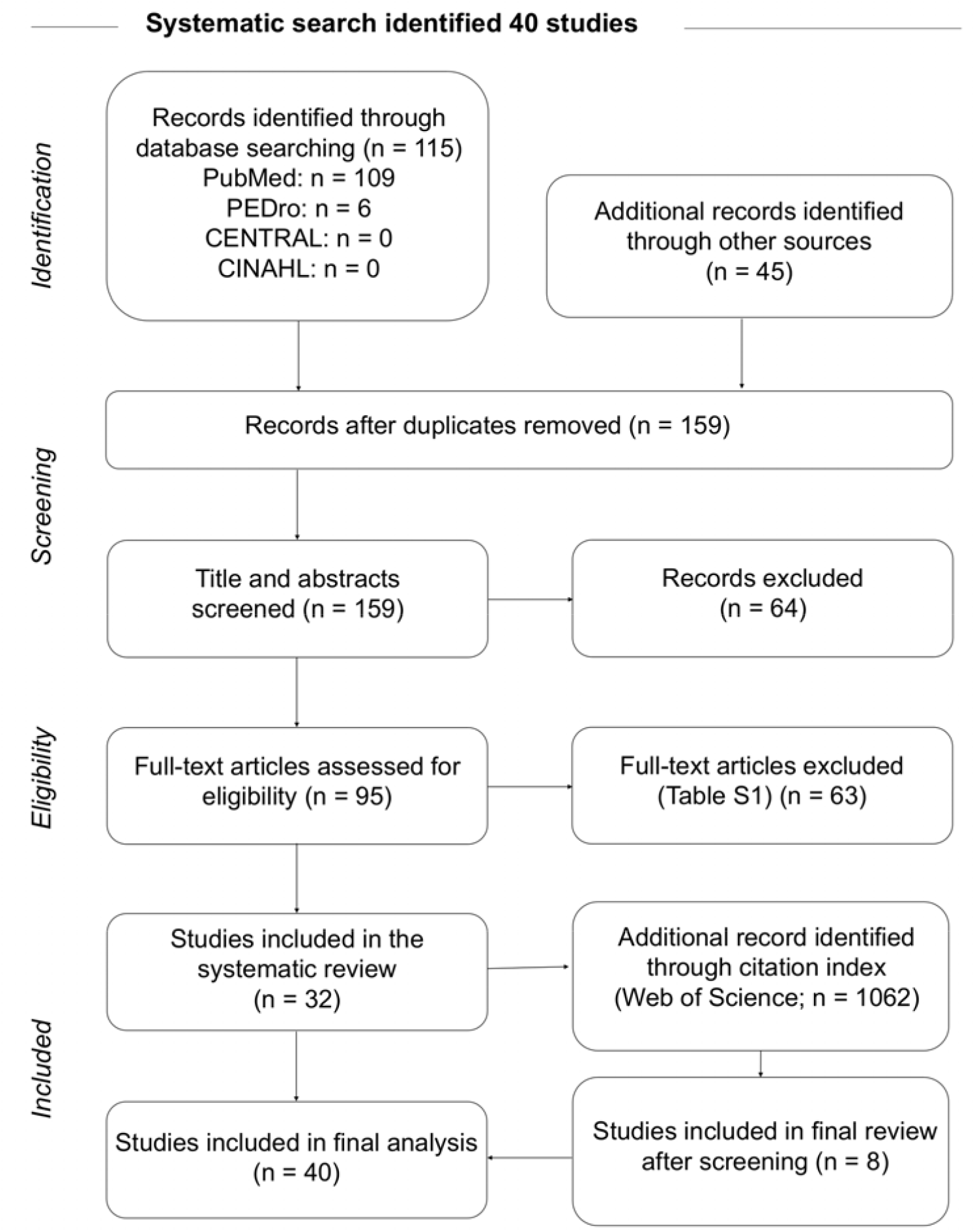
Flow diagram of literature search results. Electronic database and Google Scholar searches yielded a total of 160 studies. After duplicates were removed (n = 1), the titles and abstracts of 159 studies were screened, and the remaining 95 studies were assessed for eligibility by full-text screening. From full-text screening, 32 studies met eligibility criteria, and citation search from 1062 studies identified eight additional studies. Ultimately, a total of 40 studies were included in this study.

### Age-induced cartilage degeneration is accompanied by elevated inflammation, impaired autophagy, and cellular senescence

A meta-analysis of histological studies confirmed the expected age-related increase in cartilage degeneration, defined as a progressive loss of cartilage extracellular matrix over time (**Figure 2A and 2B**). These findings were further supported by mixed linear regression analysis (**Figure 2C**). Cartilage degeneration was also evident in middle-aged mice, consistent with clinical reports of cartilage structural abnormalities in people aged 45–55 years old(10). These gross morphological alterations were generally supported by non-pooled histology data (**Table S4**) and were also associated with lower cellularity and a higher percentage of apoptotic chondrocytes (**Figure S1**). Further supporting the evidence of age-related cartilage degeneration, aged chondrocytes secreted increased levels of MMP13 compared to young chondrocytes (**Figure S2**). MMP13 is a key enzyme in the cleavage of type II collagen (COL2A1), the predominant structural component of cartilage that is degraded in KOA, and *Mmp13* is a critical target gene in cartilage degeneration(11). Increased MMP13 expression was also associated with deposition of type X collagen (COL10A1) (**Table S4)**, suggesting aging drives chondrocytes towards a hypertrophic phenotype(12). Interestingly, one non-pooled study demonstrated that aged chondrocytes release increased levels of matrix metalloproteinase 3 (MMP3) (**Table S4**), a well-known marker of synovial inflammation in rheumatoid arthritis that is involved in the breakdown of the ECM of cartilage(13). Of note, throughout this manuscript, we use official gene symbols to refer to proteins (all caps), transcripts (sentence case), and genes (italicized), e.g. for type X collagen: COL10A1 (protein), Col10A1 (transcript), *Col10A1* (gene).

**Figure 2.**
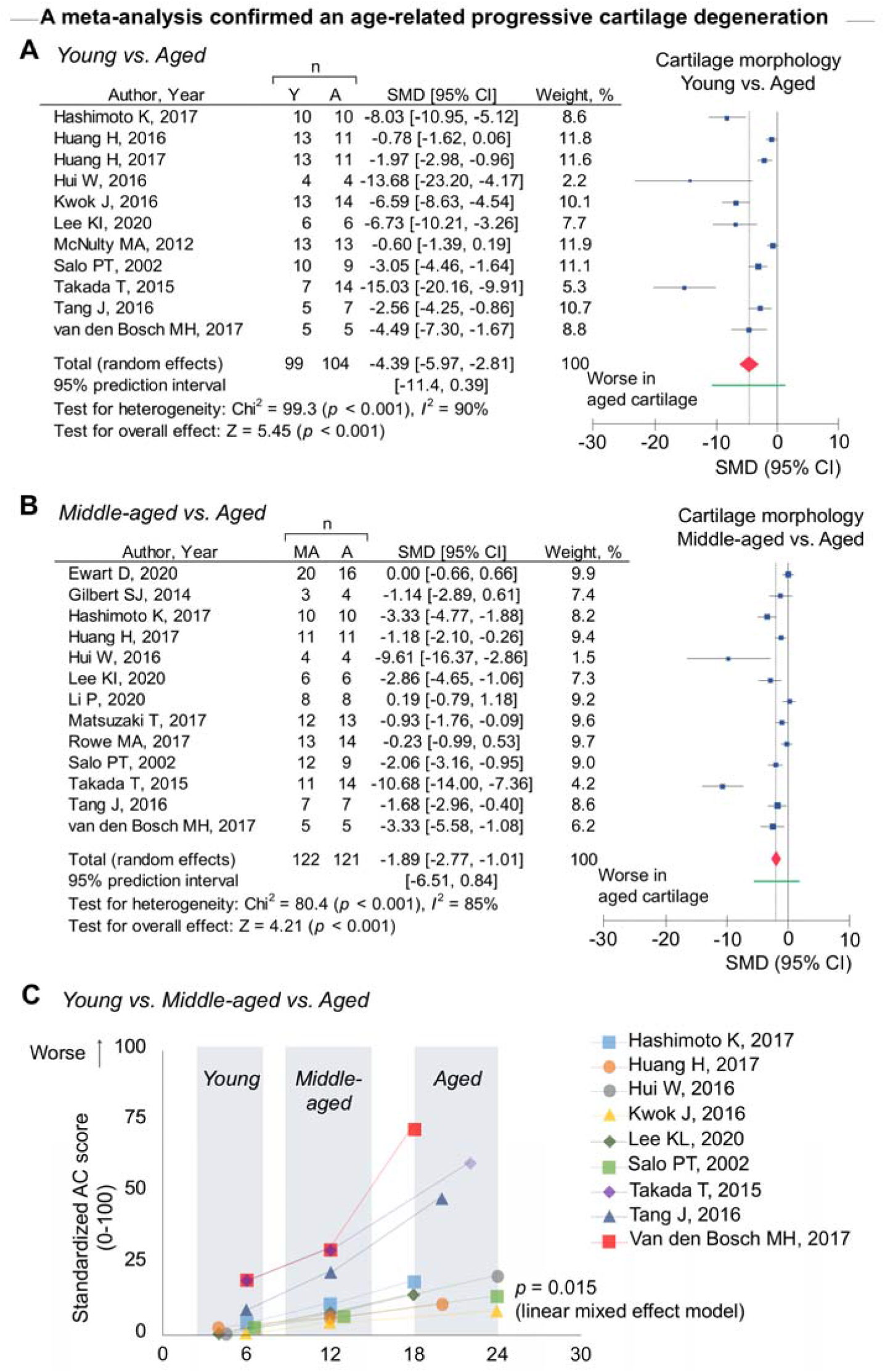
Progression of cartilage degeneration with aging. **A-B**. Effect size for cartilage morphology and age-related cartilage degeneration compared to young (A) and middle-aged cartilage (B). The forest plot displays relative weight of the individual study, standardized mean differences (SMDs), and 95% CI. The red diamond indicates the global estimate and its 95% CI. The green bar indicates the prediction interval. **C**. The trajectory of age-related alteration in cartilage morphology in individual studies. P-value of linear mixed effect model is provided.

Previous epidemiologic and biologic evidence suggest a link between age-related systemic inflammation and development of OA in humans(14). As human articular chondrocytes from older adults release higher pro-inflammatory cytokines than young counterparts(15), we probed for evidence of local inflammatory factors in aged cartilage. Although protein-level evidence of local inflammation in aged murine samples is generally lacking, one study did demonstrate that interleukin 36 receptor antagonist (IL36RA) expression was reduced with aging (**Figure S3**). IL36RA inhibits the activation of NF-κB signaling by IL-36(16), and induction of NF-κB has been associated with KOA(17). This finding suggests that the reduced IL36-RA elevates inflammation signaling, which may accelerate age-related KOA, but further investigation into protein level local inflammation is warranted.

As a next step towards a more comprehensive view of KOA, we probed for global transcriptomic cartilage changes across the lifespan of mice. Only one study to date has performed RNA-seq on cartilage tissue samples from different age groups(18). This study demonstrated that aging results in upregulation of inflammatory response-related genes and downregulation of genes associated with cartilage development and homeostasis(18). To further probe the functions of the transcript products at the pathway level, we accessed the archived RNA-seq data(18) and performed gene-set-enrichment analysis (GSEA) with summarizing redundant pathways (**Figure 3A**). The data revealed that aging upregulated inflammation associated biological processes, such as “ *antigen receptor-mediated signaling pathway*” and “ *regulation of interleukin-8 production*”, while ECM-related biological processes including “ *extracellular matrix organization*” were downregulated with aging (**Figure 3B**; see **Tables S5-S8** for details). Notably, these molecular changes were evident at middle-age (**Figure 3C**). Early changes in inflammatory processes and ECM regulation at the transcript level were further supported by a previous microarray study that showed inflammation-related genes were upregulated while ECM-related genes were downregulated in middle-aged mice(19).

**Figure 3.**
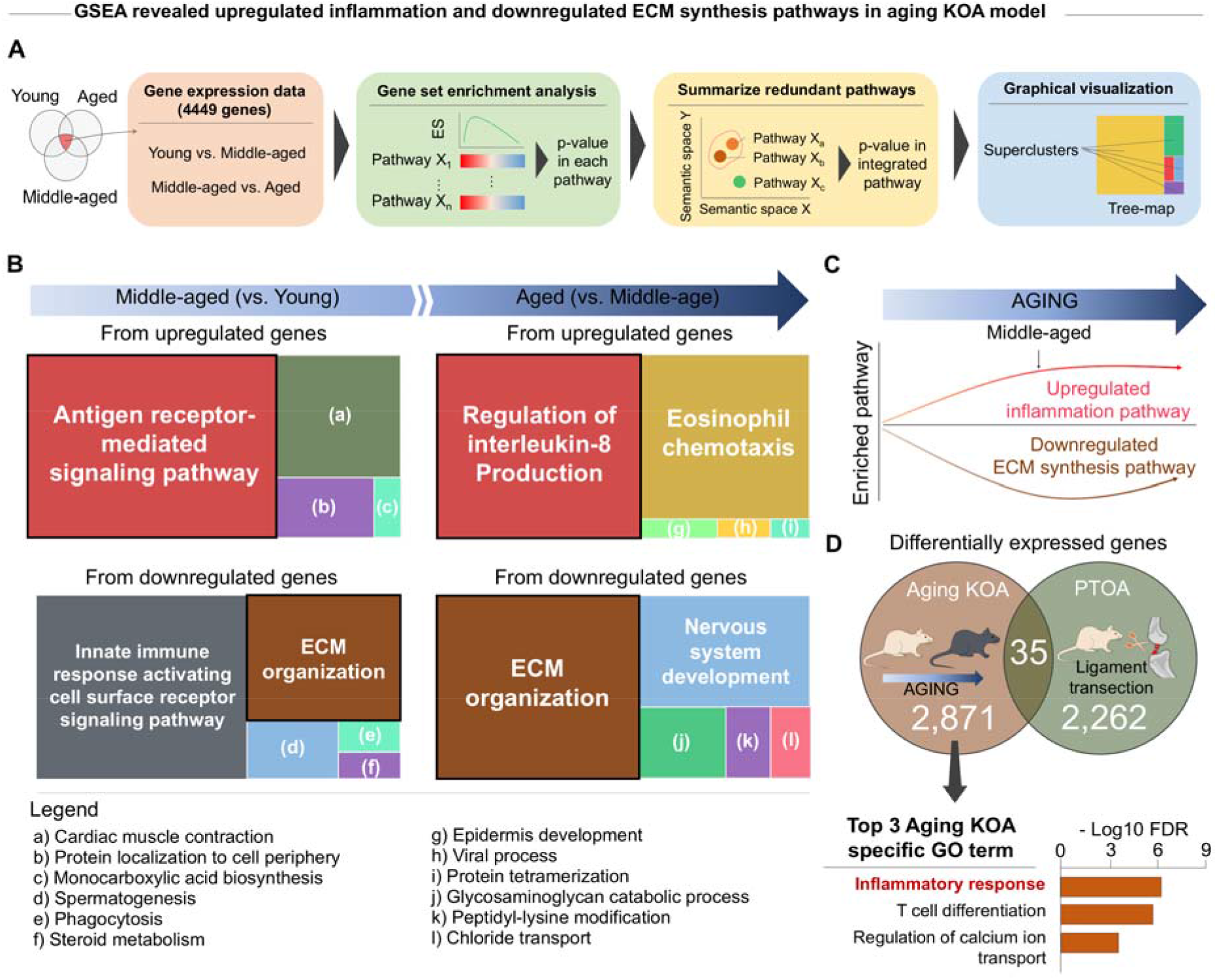
Transcriptomic signature in aged knee joints show upregulated inflammation and downregulated ECM synthesis. **A**. Analysis flow. Normalized RNA-seq data from one original study were used for gene set enrichment analysis (GSEA). REVIGO was used to summarize redundant pathway. **B**. Gene ontology (GO) enrichment tree-map of biological processes for transcripts upregulated or downregulated between young vs. middle-aged and middle-aged vs. aged. Each rectangle represents a supercluster GO, visualized with different colors. Size of rectangles was adjusted to reflect the p-value of the GO term calculated by Top GO (i.e., the larger the rectangle, the more significant the GO-term). **C**. Trajectory of inflammation and extracellular matrix (ECM) synthesis pathways predominantly identified by the GO enrichment tree-map. **D**. GO enrichment analysis from aging KOA specific genes.

To determine whether the upregulated inflammation and downregulated ECM-related pathways are unique to age-related KOA, we compared pathways between age-related KOA and PTOA models generated by differentially expressed transcripts (**Figure S4**). Surprisingly, only 35 (0.7%) of the 5,168 transcripts overlapped across the two models, suggesting that aging and PTOA have markedly distinct transcriptional profiles (**Figure 3D**). Gene ontology enrichment analysis revealed that “ *inflammatory response*” was uniquely identified in age-related KOA models. On the other hand, “ *ECM organization*” was enriched across the genes that were unique to PTOA **(Figure S4**). These data indicate distinct mechanistic trajectories when comparing PTOA versus age-related KOA, consistent with the divergent clinical phenotypes between the two pathologies(20).

To gain further insight into potential mechanisms underlying age-related cartilage degeneration, we probed cellular-level alterations as a function of time. Given the role of the inflammation-autophagy-senescence network in the pathogenesis of KOA(21), we expected to find evidence of impaired autophagy and resistance to oxidative stress, as well as increased senescence in aged chondrocytes. Indeed, impaired autophagy (as determined by decreased autophagy related 5 (ATG5)) and decreased resistance to oxidative stress (as determined by decreased heme oxygenase 1 (HMOX-1)) were observed in aged chondrocytes (**Figure S5A, B**). Although no study has directly investigated chondrocyte senescence in age-related cartilage degeneration, one study found synovial fluid in aged knee joints contained elevated senescence-related microRNAs, such as miR-34-5p (**Figure S5C**). Taken together, these findings further support the hypothesis that aging drives cartilage degeneration by modulating the integrated inflammation-autophagy-senescence network. Increased production of pro-inflammatory mediators is a prominent feature of the senescence-associated secretory phenotype (SASP)(14), which further highlights the need for evaluating the pro-inflammatory cytokines in the context of aging KOA.

### Activation of signaling molecules involved in TGF-β and AMPK signaling pathways are reduced with increasing age

We next sought to identify specific signaling pathways that contribute to age-related cartilage degeneration. Given that transforming growth factor-beta (TGF-β) protects against pro-inflammatory cytokine-mediated MMP expression in healthy joints(22) but appears to have the opposite effect in diseased joints(23), we initially focused on the TGF-β signaling pathway. Meta-analysis revealed a significant reduction in molecules involved in TGF-β signaling in aged cartilage, including TGFB2, TGFB3, phosphorylated small mothers against decapentaplegic homolog 2 (SMAD2), TGFBR1, and TGFBR2 (**Figure S6**). Activin A Receptor Like Type 1 (ACVRL1) and TGFBR1 activate SMAD1, SMAD5, & SMAD8 and SMAD2 & SMAD3, respectively(24), and genetic inhibition of *Tgfbr1* exacerbates KOA(25). Interestingly, two studies found the ratio of ACVRL1/TGFBR1 increases with age(26, 27), suggesting an age-related shift in ACVRL1/TGFBR1 may drive cartilage degeneration. Although our findings suggest downregulation of TGF-β signaling plays a role in propagating aged-related KOA, given the complexity of TGF-β signaling in KOA, its exact mechanistic role remains unclear.

Meta-analysis also revealed that aging was associated with a reduction in signaling molecules involved in 5’ AMP-activated protein kinase (AMPK) pathways, including phosphorylated protein kinase AMP-Activated catalytic subunit alpha 1 (PRKAA1), phosphorylated serine/threonine kinase 11 (STK11), proliferator-activated receptor gamma coactivator 1 alpha (PPARGC1A), and sestrin 1-3 (SESN1-3) (**Figure S7**). In light of these findings, we next probed for known transcription factors that inhibit the autophagy pathway. Forkhead box O1 (FoxO1) and FoxO3 transcription factors regulate the growth and maturation of chondrocytes(28) and have been shown in two studies to decrease with aging (**Figure S8**). Additionally, runt-related transcription factor 2 (Runx2) expression, which is partially regulated by FoxO1(29), was increased with aging (**Figure S8**). Runx2 regulates endochondral ossification and MMP13 promotor activity(30), suggesting a link between autophagy and chondrocyte hypertrophy regulation. We also found evidence that aging decreases the expression of hypoxia-inducible factor 1-alpha (Hif1a) and nuclear factor of activated T-cells, cytoplasmic 1 (Nfatc1) transcription factors (**Table S4**). Hif1a enhances ECM synthesis and regulates both autophagy and apoptosis(31), and evidence suggests Nfatc1 is essential in the cellular mechanisms driving ECM remodeling(32). These two findings provide a link between our meta-analysis and the transcript-level findings from the RNA-seq data set. Finally, transcription factor enrichment analysis from archived RNA-seq data(18) revealed that three out of five identified transcription factors (Runx2, Hif1a, and Nfatc1) were contained in the top 400 most abundant transcripts (**Table S9**). Integrating the RNA-seq data with previously reported protein-level studies supports the hypothesis that aging reduces activation of TGF-β and AMPK signaling pathways, leading to loss of protection from pro-inflammatory cytokines and ultimately, cartilage degeneration.

### Construction of protein-protein interaction networks from data across individual studies and validation by mass spectrometry reveals enrichment of AGE-RAGE signaling pathway

Complex signal transduction occurs through the dynamic modulation of protein-protein interactions (PPI), which play an important role in the development, onset, and progression of disease(33). Using a similar approach as described in the previous section, 25 differentially expressed (DE) proteins were identified from individual articles. We then probed whether these proteins are enriched into specific pathways using Kyoto Encyclopedia of Genes and Genomes (KEGG) enrichment analysis (**Figure 4A)**. As the quality of evidence of each protein was “ very low” or “ low” according to the Grades of Recommendation, Assessment, Development, and Evaluation (GRADE)(34) (**Table S10**), we also performed sensitivity analysis focusing on DE proteins with higher GRADE (i.e., high quality of evidence(35)). These two different KEGG enrichment analyses consistently identified TGF-β signaling pathway and advanced glycation end products (AGE)-receptor for AGE (RAGE) signaling pathway in diabetic complications (**Figure 4A). Figures S9-S10** contain detailed information on the risk of bias and publication bias used for GRADE assessment.

**Figure 4.**
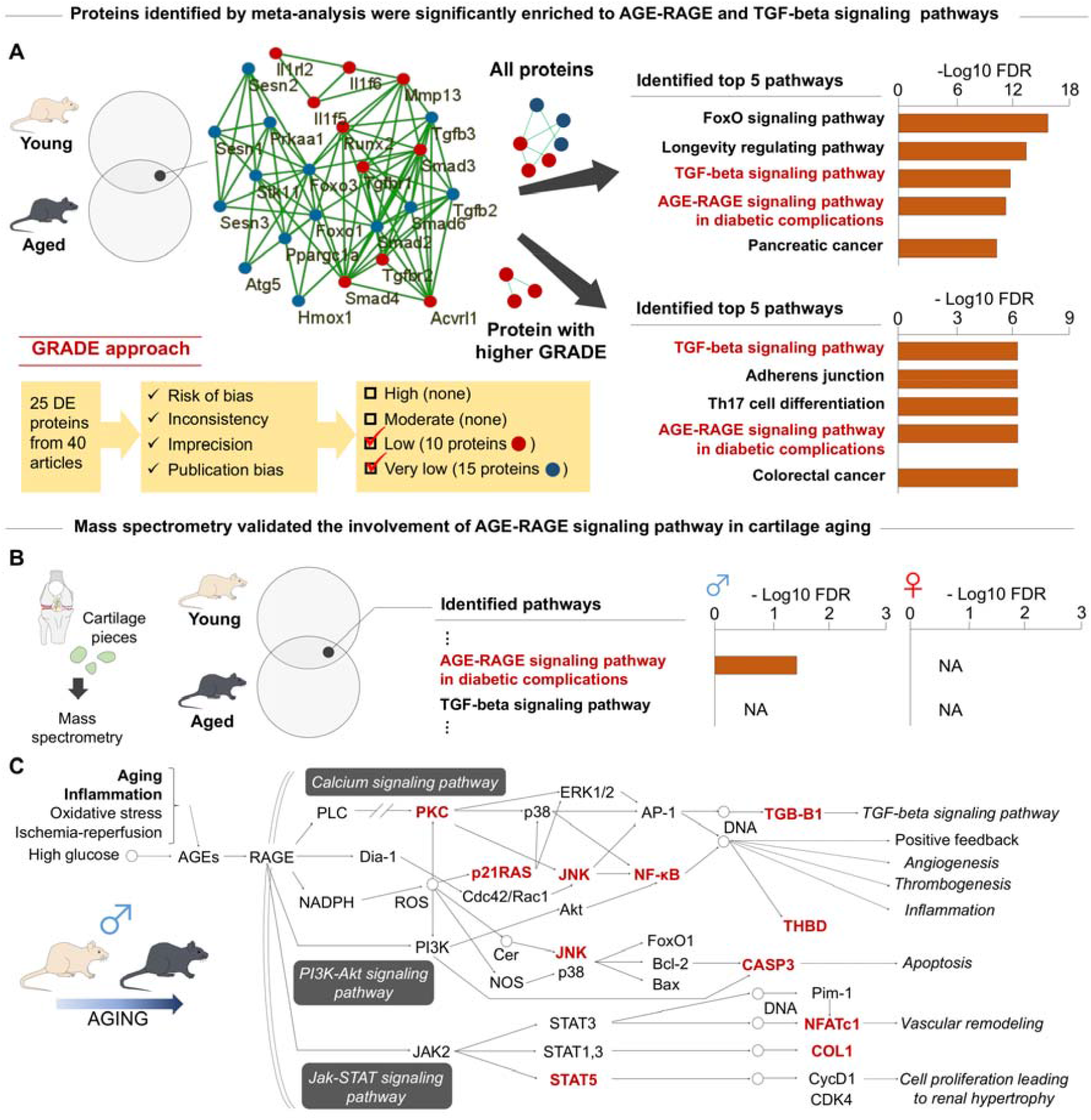
AGE-RAGE signaling pathway involved in cartilage aging. **A**. KEGG enrichment analysis to identify specific pathways associated with cartilage aging based on 25 differentially expressed (DE) proteins (p<0.05) between young vs. aged identify by previous articles. As a sensitivity analysis, we included 10 proteins of higher the Grades of Recommendation, Assessment, Development, and Evaluation (GRADE) score. Interactome network was generated by STRINGdb for DE proteins between young vs. aged. Each node represents a protein identified from meta-analysis and each line represents an interaction. Line thickness reflects the strength of the interaction. Color of nodes refers to the GRADE score; red represents low GRADE and blue represents very low GRADE. **B**. Mass spectrometry of the DE proteins between young vs. aged C57/BL6 mice to validate the two pathways identified by meta-analysis (i.e., TGF-beta and AGE-RAGE signaling pathways). We then performed KEGG enrichment analysis and identified AGE-RAGE signaling pathway in male mice. **C**. Visualization of AGE-RAGE signaling pathway. Each node represents a protein, and the red node represents the DE proteins (p<0.05) confirmed with cartilage aging in male mice.

To validate the involvement of the two pathways identified by meta-analysis, we performed mass spectrometry proteomic analysis of data from a recently completed study that included cartilage tissue from young (4 months old) and aged (21 months old) C57/BL6 male and female mice (**Figure 4B**)(36). KEGG enrichment analysis from the mass spectrometry data confirmed AGE-RAGE signaling pathway in diabetic complications was upregulated with increasing age (**Figure 4B, C**). Most notably, the change in AGE-RAGE signaling over time was exclusively observed in male mice, thereby highlighting the importance of considering sex as a biological variable in clarifying the pathogenesis of age-induced KOA. Although AGE-RAGE signaling is a well-known signaling cascade in diabetes(37), these results indicate that AGE-RAGE signaling pathway may also be involved in age- and sex-dependent cartilage degeneration.

### Increased transparency in reporting outcomes is imperative for improving the translatability of findings in KOA research

While transparent reporting is an essential criterion for any research endeavor, it is all-too-often overlooked(38). To encourage continued efforts to increase rigor in the field, we evaluated the reporting quality of the 40 studies included in this study using ARRIVE guidelines(8) (**Figure S11A**). Among the most pervasive weaknesses, we found that none of the studies adequately reported experimental outcomes (item 12) or adverse events (item 17), and most studies lacked detailed information describing experimental animals including sex (item 8), housing and husbandry (item 9), sample size (item 10), and baseline data (item 14). The lack of such information makes it impossible to reliably reproduce the experimental findings and generalize these studies to broader populations. Importantly, approximately 30% of the articles included in the meta-analysis lacked information regarding sex, and none of the studies considered sex as a biological variable. As mentioned above, mass spectrometry data indicate that signaling pathways associated with cartilage aging are influenced by sex, further supporting the need for consideration of sex as a biological variable.

Though the current state of reporting is still less than optimal, the field is showing an encouraging trend overall (**Figure S11B**). After the publication of the ARRIVE guidelines in 2010, studies displayed a significantly higher quality of reporting compared to those published prior to 2010 (**Figure S11C**). However, there is a continued and urgent need for increased attention to steps ensuring rigor and reproducibility in studies in order to accelerate the translation of interventions designed to treat age-related KOA and improve patient care, especially in female.

## DISCUSSION

Using an integrated approach of meta-analysis and multi-omics analyses, this study synthesized age-related changes in murine knee joint articular cartilage based on available evidence from 40 different murine studies and supporting proteomic analysis (**Figure 5**). By excluding surgically induced PTOA models, our findings provide novel insights into the unique pathways associated with aging, the most common risk factor for KOA by far. Consistent with clinical reports(10), meta-analysis revealed that age-related cartilage degeneration was apparent in middle-aged mice both histologically and at the transcription level. The onset of the impaired cartilage integrity was accompanied by upregulation of genes associated with inflammation, which is a prominent feature in age-related KOA. Construction of protein-protein interaction networks from data across individual studies revealed enrichment of AGE-RAGE signaling pathway and was validated by mass spectrometry. As inflammation is a potential trigger of this pathway in cartilage, our findings highlight the need for future studies to further probe the mechanistic role of inflammation and AGE-RAGE signaling pathway on age-related KOA.

**Figure 5.**
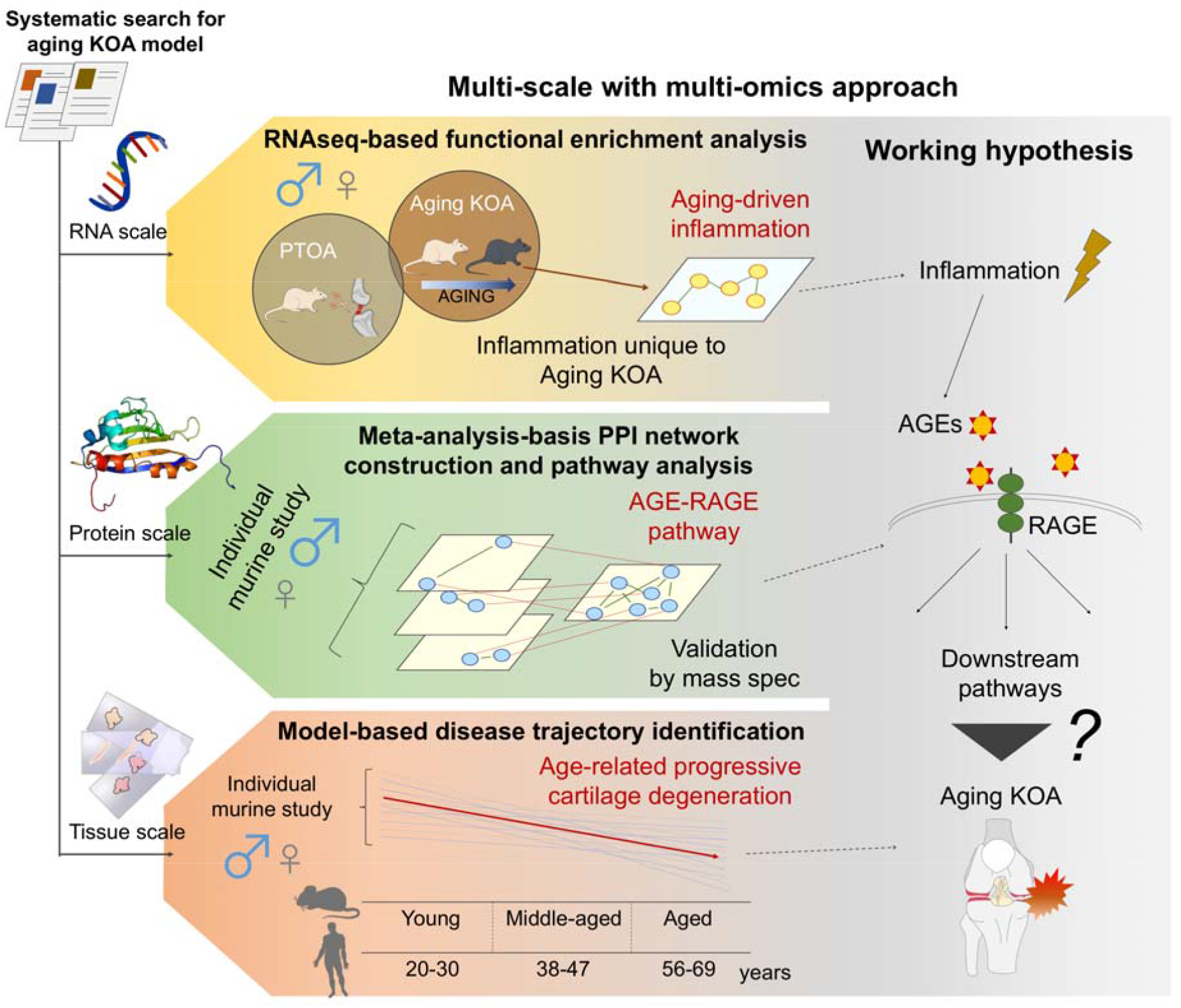
Graphical abstract. We appraised and collated data from 40 available murine studies to provide insight into unique pathways associated with age-related KOA using multi-scale (RNA, protein, and tissue scales) with multi-omics approach. Meta-analysis revealed that age-related progressive cartilage degeneration was apparent in middle-aged mice (tissue scale), which is accompanied by transcriptomic signature of aging-driven upregulated inflammation (RNA scale). Our approach of bioinformatics with meta-analysis of aging-associated proteins revealed altered AGE-RAGE signaling as a key driver of age-related KOA (protein scale). Most notably, mass spectrometry revealed the AGE-RAGE signaling pathway to be identified only in male mice, highlighting the importance of considering sex as a biological variable in clarifying the pathogenesis of age-related KOA. This evidence indicates that AGE-RAGE signaling pathway may be uniquely involved in age- and sex-dependent cartilage degeneration with the elevated inflammatory response.

Animal studies typically focus on individual proteins or pathways, thereby precluding a holistic understanding of disease pathogenesis. Here, we implemented a network-based approach to identify possible pathways associated with age-related KOA. We found that AGE-RAGE signaling was the only significantly enriched pathway with increasing age. Evidence for the involvement of the AGE-RAGE pathway in murine knee cartilage supports the accumulation of AGE observed in aged human knee cartilage(39). Accumulated AGE suppresses proteoglycan synthesis in mouse chondrocytes(40) and increases MMP13 release in human chondrocytes(41), a hallmark of age-related cartilage degeneration that was confirmed by our meta-analysis. Although the signaling cascade involved in the AGE-RAGE pathway is not fully understood in the context of KOA, RAGE signaling activates inflammatory pathways and increases MMP-13 expression in human chondrocytes(42). Furthermore, AGE increases the stiffness of the collagen network in human knee cartilage(39), altering mechanotransductive-dependent chondrogenesis pathways(43). The evidence from murine and human studies indicates that limiting AGE accumulation may be a promising therapeutic target to prevent or slow the progression of age-related KOA. Despite the demonstrated role of AGE accumulation in cartilage aging, the mechanisms leading to AGE synthesis and accumulation in aged cartilage remain poorly understood. This study highlights age-related inflammation as a potential trigger for AGE accumulation in cartilage, thereby representing an important topic of future research.

In addition to the mechanistic insights gained through integrative meta-analysis and a systems biology approach, our work also highlights the need for study designs that ensure methodological rigor and consider sex as a biological variable. Despite the fact that women have a greater risk of developing KOA than men(9), the overwhelming majority of studies investigating KOA only included male mice. Yet, there are currently no published studies that compared KOA pathogenesis according to sex. Sex as a biological variable is clearly an essential consideration as the field strives to increase rigor, promote discovery, and expand the clinical relevance of age-related KOA murine models. Indeed, mass spectrometry-based proteomic analysis identified the AGE-RAGE signaling pathway only in male mice, further highlighting the need for the consideration of sex in cartilage aging research.

Although this study provides a new perspective to the pathogenesis of KOA, it has limitations. Whereas our study only focused on the cartilage of the knees, it is well established that KOA is a disease of the whole joint. Additionally, given the lack of female mice in current studies, we were not able to probe mechanisms underlying age-related KOA in female mice. This remains an important area for future work.

According to both histological observation and transcriptomic analysis, our findings suggest that the pathogenesis of KOA in natural aging murine models follows a similar relative timeline to that observed in humans. Moreover, we found that declines may be driven initially by increased inflammation. Our results further revealed that the AGE-RAGE signaling pathway may be uniquely involved in age-related cartilage degeneration, linking the observed changes in ECM with the intense inflammatory response. In pre-clinical models, increased attention should be paid to the differences between age-associated pathology and trauma-related pathology, as our findings suggest that distinct mechanisms between the two exist. These findings are in line with subgrouping amongst clinical phenotypes(20). Finally, we identified several shortcomings of the current state-of-the-science, including a lack of transparent reporting and a lack of consideration of sex as a biological variable. Together, these barriers likely contribute to delays in translational efforts. This work highlights the need for scientists to adopt targeted steps to enhance scientific rigor in the field in order to accelerate the pace of discovery toward the development of safe and effective KOA interventions.

## MATERIALS AND METHODS

The full methodology for this manuscript is provided in detail in the supplemental file and includes information on: Eligibility Criteria, Literature Search, Study Selection, Data Collection, Meta-Analysis, Functional Characteristics of Transcriptome, Protein-Protein Interaction Network, Reporting Quality (ARRIVE), Risk of Bias (SYRCLE), Publication Bias, and Quality of Evidence (GRADE).

### Literature search, study selection, and data collection

We included articles characterizing aged articular cartilage in the knee joints of mice (i.e., each study had to include both aged and young mice or aged and middle-aged mice). Young, middle-aged, and aged mice were defined as 2-7, 9-15, and 18-24 months old, respectively, and correspond to approximately ages 20-30, 38-47, and 56-69 years in humans, respectively(44). An electronic search was conducted on March 10, 2020. A manual search of the reference lists within previously published narrative reviews was also performed. Finally, a citation search was performed on the original records with Web of Science. Two independent reviewers (HI and GG) assessed eligibility and screened titles and abstracts yielded by the search. Full manuscripts of the articles that met the eligibility criteria were then reviewed. Disagreements regarding manuscript inclusion between the two reviewers was discussed until consensus was achieved.

A single reviewer (HI) extracted data regarding basic study information (authors, publication year, and country of the corresponding author), experimental condition (i.e., mice strain, age, sample size, and sex), target joint (tibiofemoral or patellofemoral joints), outcome measures (cartilage morphology, morphometry, composition, biomechanical characterization, biomarker, and molecular biology), funding, and presence of conflict of interests.

### Meta-analysis

To characterize aged articular cartilage and the underlying mechanism of age-related cartilage degeneration, pooled estimates and 95% confidence intervals for standardized mean differences (SMD) of outcome measures were calculated using a random-effect model. SMD were calculated using the mean between-group difference (aged and middle-aged or young) divided by the pooled standard deviation. Study heterogeneity, the inter-trial variation in study outcomes, was assessed using *I*^*2*^, which is the proportion of total variance explained by inter-trial heterogeneity.

To address the trajectory of age-related changes in articular cartilage, a mixed linear regression analysis with random slopes and random intercepts was performed for the outcome of cartilage morphology. In this analysis, age category (1: young, 2: middle-aged, 3: aged) and standardized semi-quantitative score of cartilage degeneration was included as independent and dependent variables, respectively. To standardize semi-quantitative score of cartilage degeneration, all histological scores provided in each included study were converted to 0-100 and recalculated as in a previous meta-analysis(45), with higher score indicates severe cartilage degeneration.

### Multi-omics computational analysis

To evaluate function of transcripts that were significantly altered with cartilage aging, we performed single sample gene set enrichment analysis (ssGSEA). R/Bioconductor package fgsea was used with including gene scores defined by log2 fold change of gene expression profile available from one study(18). This analysis can determine whether a given gene set is significantly enriched in a list of gene markers ranked by their correlation with a phenotype of interest. We used GO terms (GO_Biological_Process_2018) as a gene set downloaded from enrichr (https://amp.pharm.mssm.edu/Enrichr/). Subsequently, REVIGO software(46) was applied to summarized redundant GO terms and visualize the summarized results.

Next, with the goal of identifying transcription factors that regulate target genes associated with cartilage aging, we performed transcription factor enrichment analysis using ChIP-X Enrichment Analysis Version 3 (ChEA3)(47). RNA-seq data (18) with false discovery rate adjusted p-value less than 0.05 was used.

We then used the Search Tool for the Retrieval of Interacting Genes database (STRINGdb)(48) as a means to evaluate the PPI across proteins that were significantly associated with cartilage aging. Kyoto Encyclopedia of Genes and Genomes (KEGG) pathway enrichment analyses using the STRINGdb(48) was then performed in order to determine whether proteins associated with cartilage aging were significantly enriched in specific signaling pathways. In these analyses, we included significantly expressed proteins identified by the meta-analysis or mass-spectrometry proteomics available from one study(36).

### Overall quality of evidence

Two independent reviewers (HI and KW) assessed the reporting quality (ARRIVE)(8), risk of bias, publication bias, and quality of evidence (GRADE)(34). Disagreements between the two reviewers were discussed until consensus was achieved. For GRADE assessment, The evidence quality was downgraded if (1) outcomes have a high risk of bias; we defined this as a lack of blinding outcome assessment in more than 50% of the included studies (risk of bias domain); (2) heterogeneity between trials was more than substantial (*I*^*2*^ ≥ 50%) with non-overlapping 95% CI (inconsistency domain); (3) sample size was inadequate; we defined this as optimal information size(49) and wide 95% CI that included 0 (precision domain); and (4) publication bias existed, as identified by the Egger’s regression test (publication bias domain). The publication bias domain was applied only for outcome with ≥10 studies. Downgrade in the indirectness domain was not applied, as all studies used mouse models and outcomes were not directly related to human clinical trials and clinical decisions. The quality of evidence was judged as “ high”, “ moderate”, “ low,” or “ very low”.

## Acknowledgements

This study was supported in part by (1) a Grant-in-Aid from the Japan Society for the Promotion of Science for Overseas Research Fellowships for HI (grant no. N/A), (2) NIA R01AG052978 for FA, and (3) the National Institute of General Medical Sciences of the National Institutes of Health under Award Number T32GM008208 for GG. The funders had no role in study design, data collection and analysis, decision to publish, or preparation of the manuscript.

## Author contributions

All authors made substantial contributions in the following areas: (1) conception and design of the study, acquisition of data, analysis and interpretation of data, drafting of the article; (2) final approval of the article version to be submitted; and (3) agreement to be personally accountable for the author’s own contributions and to ensure that questions related to the accuracy are appropriately investigated, resolved, and the resolution documented in the literature.

The specific contributions of the authors are as follows:

H.I., G.G., Y.M., F.A. provided the concept, idea and experimental design for the studies. H.I., G.G., and F.A. wrote the manuscript. H.I., G.G., K.W., S.S., C.E., Y.M., and F.A. provided data collection, analyses, interpretation and review of the manuscript. H.I., G.G., and F.A. obtained funding for the studies.

