## Supplemental Appendix S1 for "Meta-analysis and multi-omics to elucidate pathogenic mechanisms of age-related knee osteoarthritis"

**The PDF file includes:**

Figure S1. Age-related change in anti-apoptotic activity or apoptosis (BCL2, BAG1)

Figure S2. Age-related change in ECM degradation enzyme (MMP13)

Figure S3. Age-related change in pro-inflammatory cytokine (IL36A, IL1RL2, IL36RA)

Figure S4. Figure S4. Functional characterization of transcriptome in aging KOA and PTOA models

Figure S5. Age-related change in autophagy, anti-oxidative ability, and senescence (LC3, Nitrotyrosine, HMOX1, miR146a-5p, miR128a-3p, miR146a-5p)

Figure S6. Age-related change in TGF-beta signaling (TGFB-1, TGFB-2, TGFB-3, SMAD2, SMAD3, SMAD4, SMAD6, SMAD7, ACVRL1, TGFBR1, TGFBR2, SMAD2P)

Figure S7. Age-related change in AMPK signaling (PRKAA1p, pSTK11, PPARGC1A, SESN1, SESN2, SESN3)

Figure S8. Age-related change in transcription factor (FoxO1, FoxO3, Runx2)

Figure S9. Risk of bias assessment using SYRCLE

Figure S10. Publication bias for cartilage morphology

Figure S11. Reporting quality of individual studies using ARRIVE guideline

Supplementary methods

Supplemental references

This supplementary material has been provided by the authors to give readers additional information about their work.

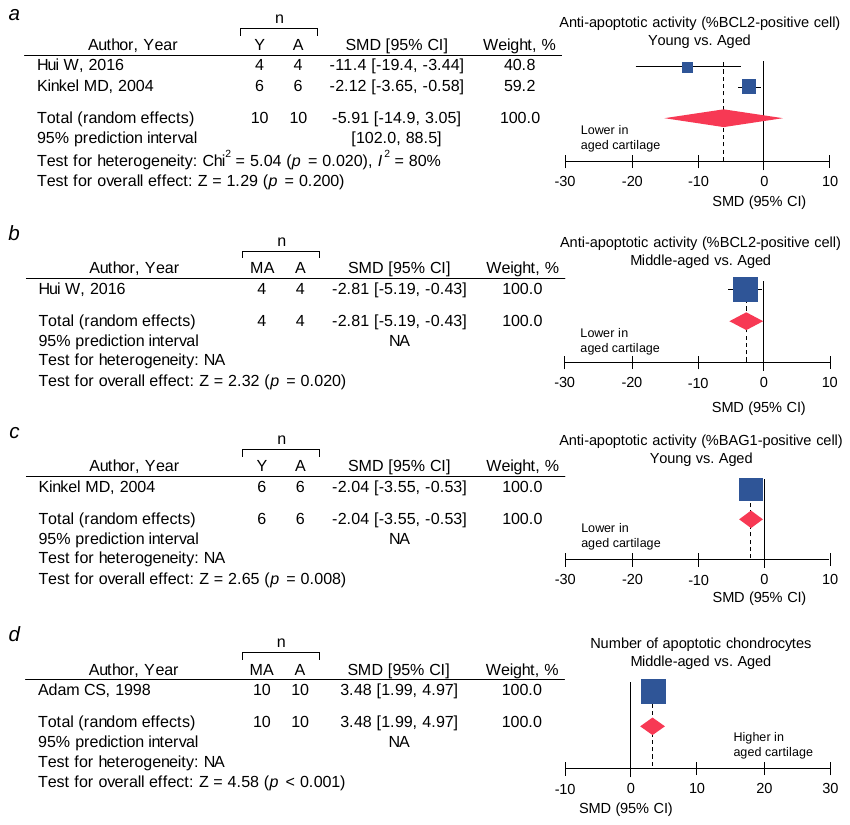

**Figure S1. Age-related change in anti-apoptotic activity and apoptosis**

***a*.** %BCL2-positive chondrocytes (young vs. aged). ***b*.** %BCL2-positive chondrocytes (middle-aged vs. aged). ***c*.** %BAG1-positive chondrocytes (young vs. aged). ***d.*** Number of apoptotic chondrocytes (middle-aged vs. aged). The red diamond represents the pooled effect size (standardized mean difference; SMD). The vertical solid line at 0 represents no difference. The prediction interval is not displayed because of wide range. Y: young, MA: middle-aged, Y: aged.

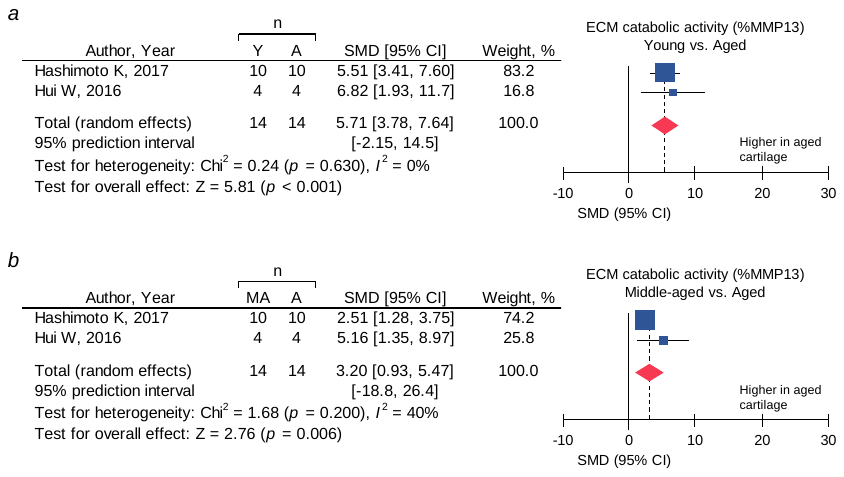

**Figure S2. Age-related change in ECM degradation enzyme**

***a*.** %MMP13-positive chondrocytes (young vs. aged). ***b*.** %MMP13-positive chondrocytes (middle-aged vs. aged). The red diamond represents the pooled effect size (standardized mean difference; SMD). The vertical solid line at 0 represents no difference. The prediction interval is not displayed because of wide range. Y: young, MA: middle-aged, Y: aged.

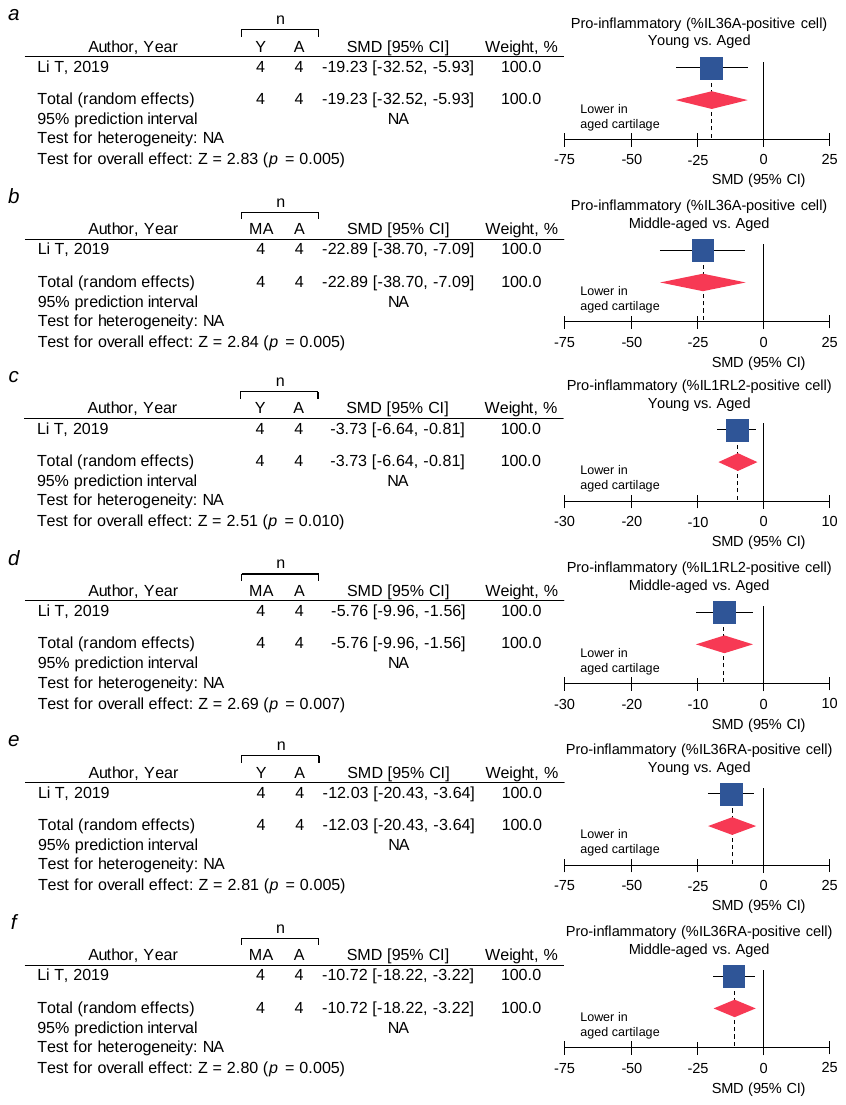

**Figure S3. Age-related change in pro-inflammatory cytokine**

***a.*** %IL36A-positive chondrocytes (young vs. aged). ***b*.** %IL36A-positive chondrocytes (middle-aged vs. aged). ***c*.** %IL1RL2-positive chondrocytes (young vs. aged). ***d*.** %IL1RL2-positive chondrocytes (middle-aged vs. aged). ***e*.** %IL36RA-positive chondrocytes (young vs. aged). ***f*.** %IL36RA-positive chondrocytes (middle-aged vs. aged). The red diamond represents the pooled effect size (standardized mean difference; SMD). The vertical solid line at 0 represents no difference. Y: young, MA: middle-aged, Y: aged.

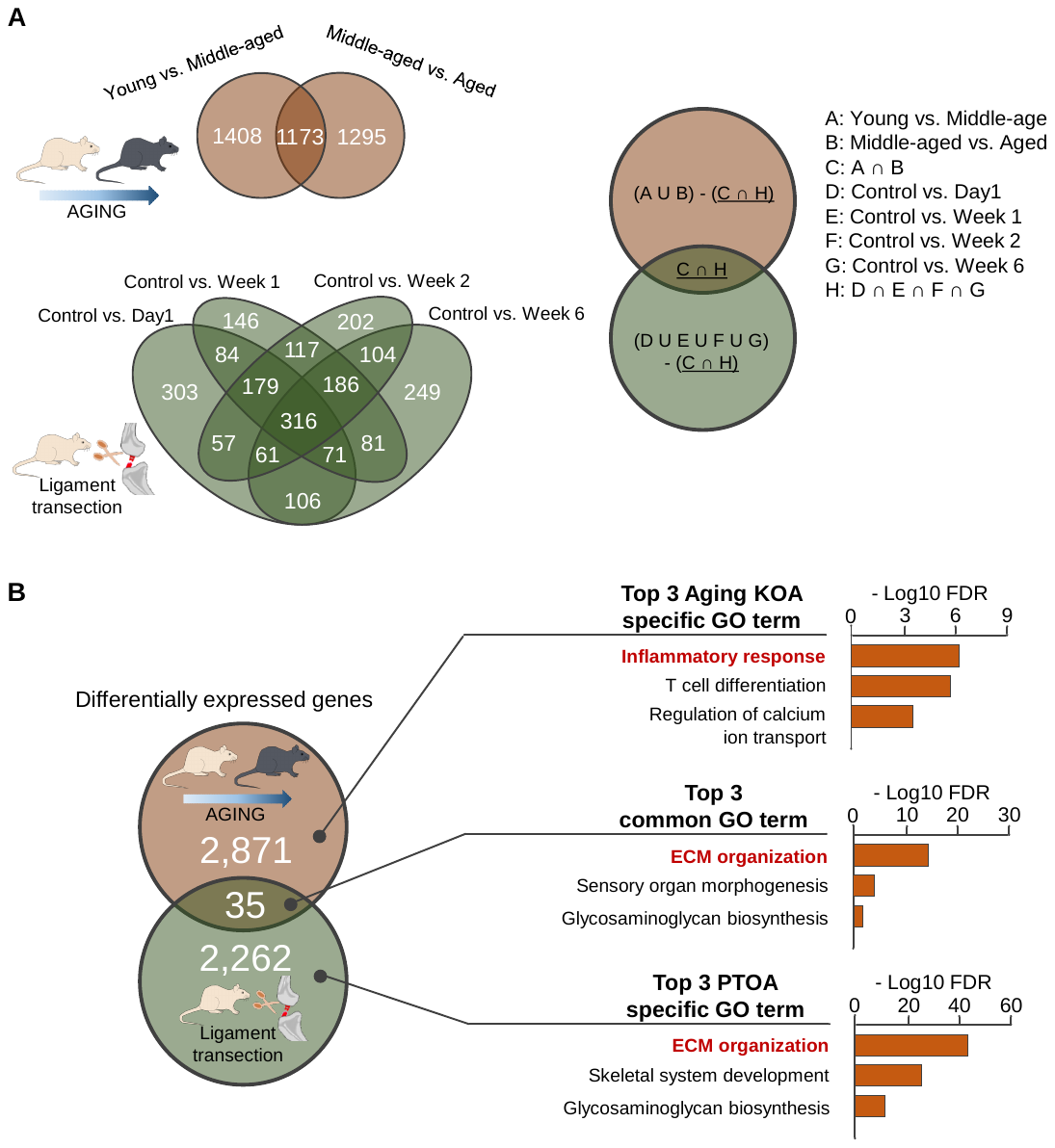

**Figure S4. Functional characterization of transcriptome in aging KOA and PTOA models**

**A.** Definition of aging KOA specific genes, PTOA specific genes, and common genes between the two models. We accessed the archived RNA-seq data from aging KOA (young vs. middle-aged, middle-aged vs, aged), and surgically induced PTOA models (control vs. day 1, control vs. week 1, control vs. week 2, and control vs. week 6). Value in the Venn diagram indicates number of genes in each category.

**B.** GO enrichment analysis for transcripts in aging KOA specific (2,871 genes), PTOA specific (2,262 genes) and common genes across two models (35 genes). GO enrichment analysis was performed by Enrichr software. REVIGO was used to summarize redundant pathway. Bar graph illustrates top 3 GO terms in each group.

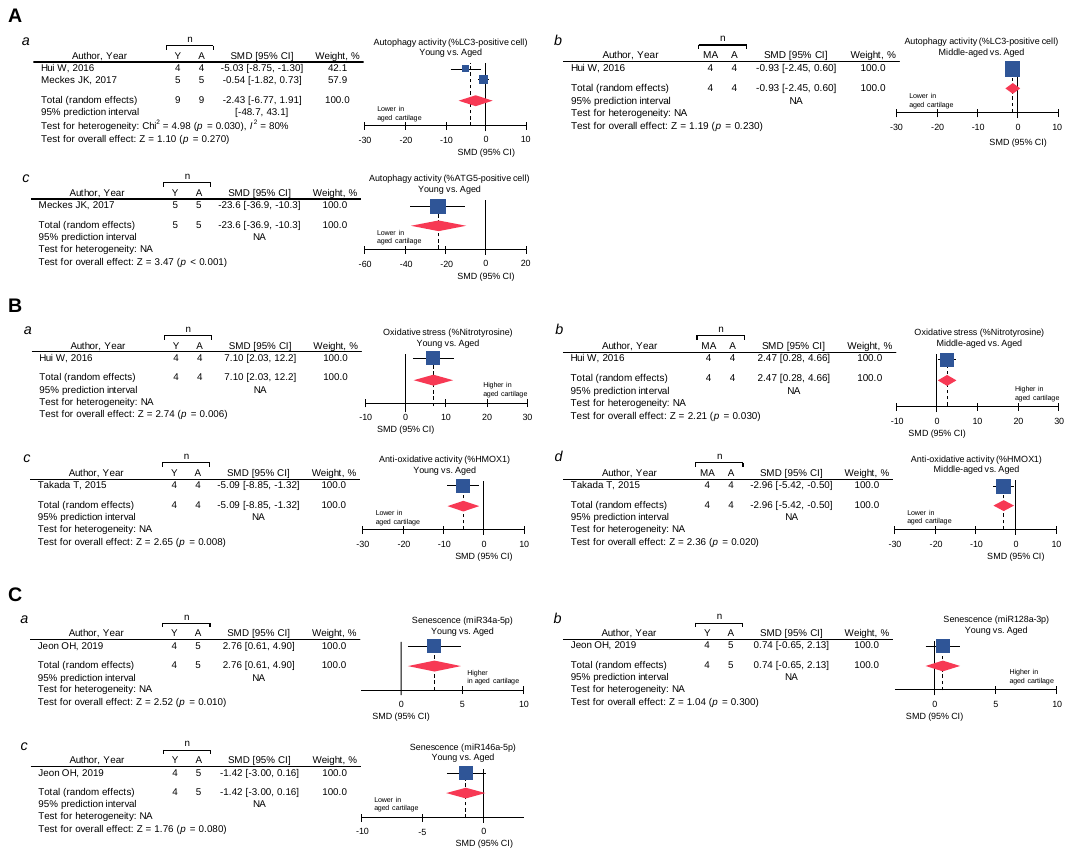
**Figure S5. Age-related change in autophagy, anti-oxidative ability, and senescence**

**A.** Between-group difference in autophagy. ***a***. %LC3-positive chondrocytes (young vs. aged). ***b*.** %LC3-positive chondrocytes (middle-aged vs. aged). ***c*.** %ATG5-positive chondrocytes (young vs. aged). **B.** Between-group difference in anti-oxidative ability. ***A*.** %Nitrotyrosine-positive chondrocytes (young vs. aged). ***b*.** %Nitrotyrosine-positive chondrocytes (middle-aged vs. aged). ***c*.** %HMOX1-positive chondrocytes (young vs. aged). ***d*.** %HMOX1-positive chondrocytes (middle-aged vs. aged).

**C.** Between-group difference in senescence. ***a*.** miR34a-5p (young vs. aged). ***b*.** miR128a-3p (young vs. aged). ***c*.** miR146a-5p (young vs. aged). The red diamond represents the pooled effect size (standardized mean difference; SMD). The vertical solid line at 0 represents no difference. The prediction interval is not displayed because of wide range. Y: young, MA: middle-aged, Y: aged.

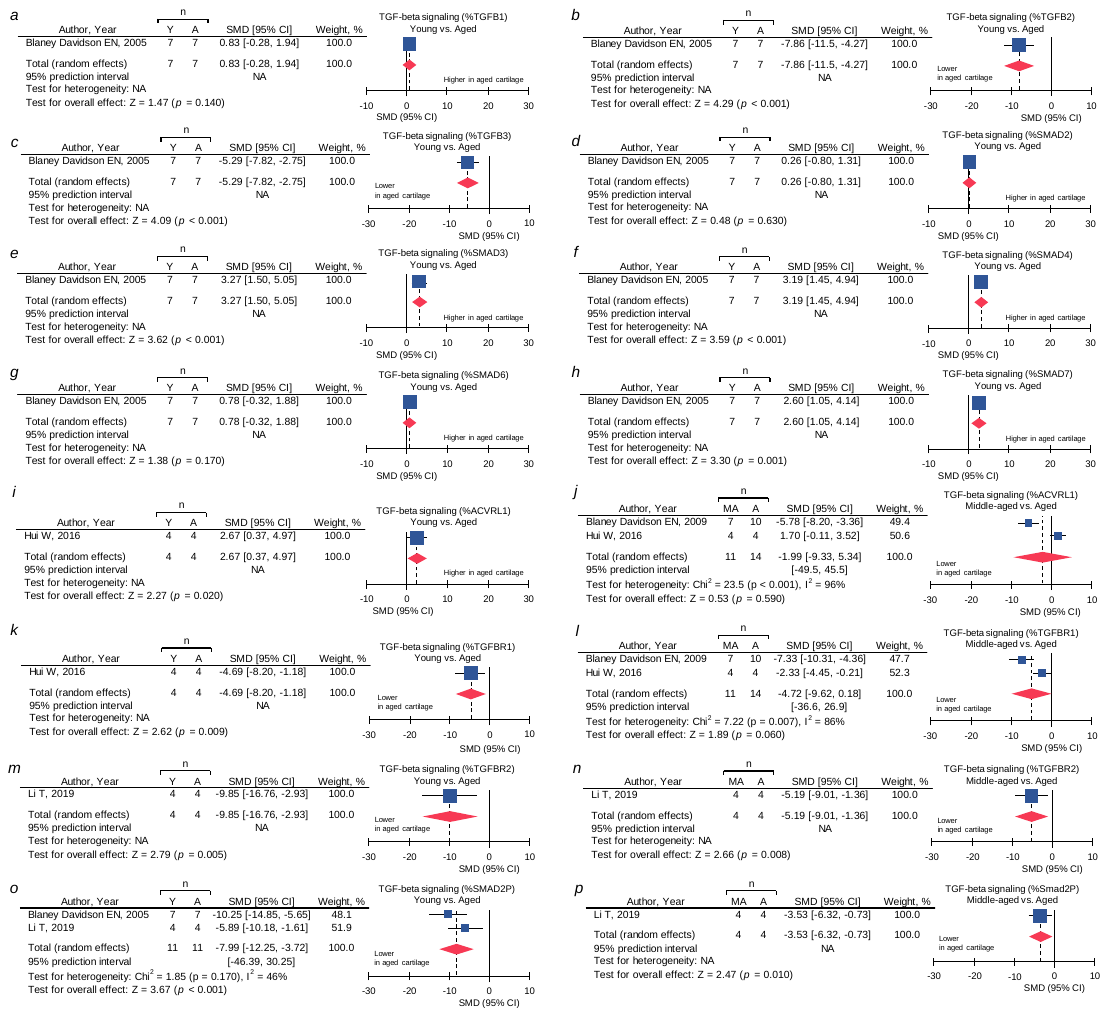

**Figure S6. Age-related change in TGF-beta signaling**

***a*.** %TGFB1-positive chondrocytes (young vs. aged). ***b*.** %TGFB2-positive chondrocytes (young vs. aged). ***c*.** %TGFB3-positive chondrocytes (young vs. aged). ***d*.** %SMAD2-positive chondrocytes (young vs. aged). ***e*.** %SMAD3-positive chondrocytes (young vs. aged). ***f*.** %SMAD4-positive chondrocytes (young vs. aged). ***g*.** %SMAD6-positive chondrocytes (young vs. aged). ***h*.** %SMAD7-positive chondrocytes (young vs. aged). ***i*.** %ACVRL1-positive chondrocytes (young vs. aged). ***j*.** %ACVRL1-positive chondrocytes (middle-aged vs. aged). ***k*.** %TGFBR1-positive chondrocytes (young vs. aged). ***l*.** %TGFBR1-positive chondrocytes (middle-aged vs. aged). ***m*.** %TGFBR2-positive chondrocytes (young vs. aged). ***n*.** %TGFBR2-positive chondrocytes (middle-aged vs. aged). ***o*.** %SMAD2P-positive chondrocytes (young vs. aged). ***p*.** %SMAD2P-positive chondrocytes (middle-aged vs. aged). The red diamond represents the pooled effect size (standardized mean difference; SMD). The vertical solid line at 0 represents no difference. The prediction interval is not displayed because of wide range. Y: young, MA: middle-aged, Y: aged.

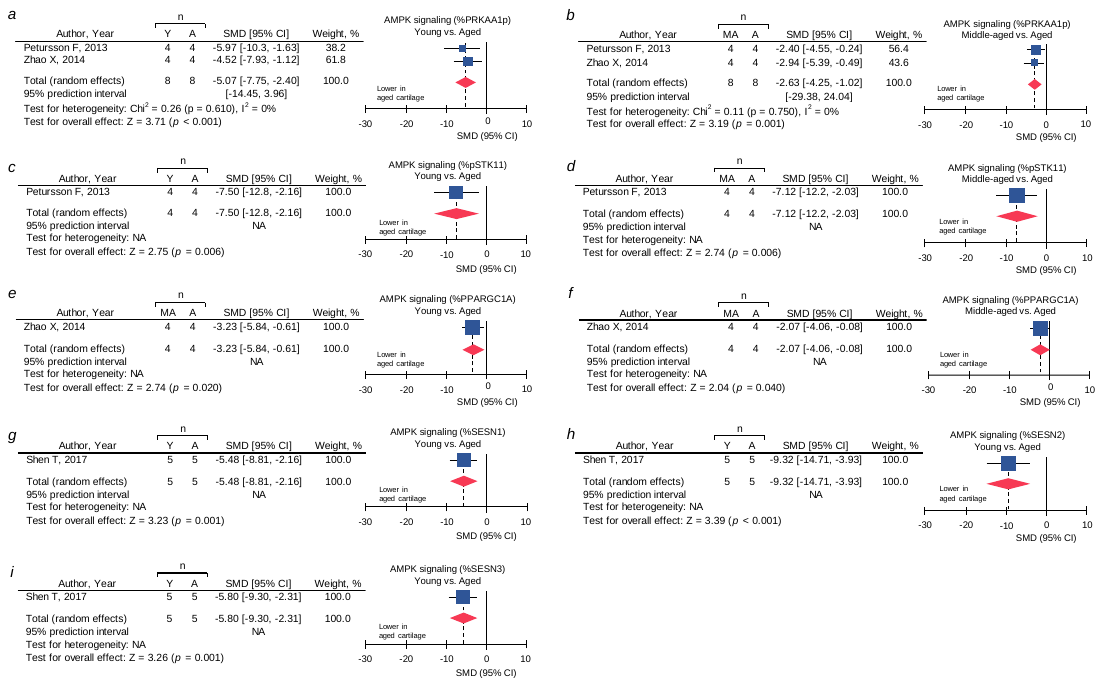

**Figure S7. Age-related change in AMPK signaling**

***a*.** %PRKAA1p-positive chondrocytes (young vs. aged). ***b*.** %PRKAA1p-positive chondrocytes (middle-aged vs. aged). ***c*.** %pSTK11-positive chondrocytes (young vs. aged). ***d*.** %pSTK11-positive chondrocytes (middle-aged vs. aged). ***e*.** %PPARGC1A-positive chondrocytes (young vs. aged). ***f*.** %PPARGC1A-positive chondrocytes (middle-aged vs. aged). ***g*.** %SESN1-positive chondrocytes (young vs. aged). ***h*.** %SESN2-positive chondrocytes (young vs. aged). ***i*.** %SESN3-positive chondrocytes (young vs. aged). The red diamond represents the pooled effect size (standardized mean difference; SMD). The vertical solid line at 0 represents no difference. The prediction interval is not displayed because of wide range. Y: young, MA: middle-aged, Y: aged.

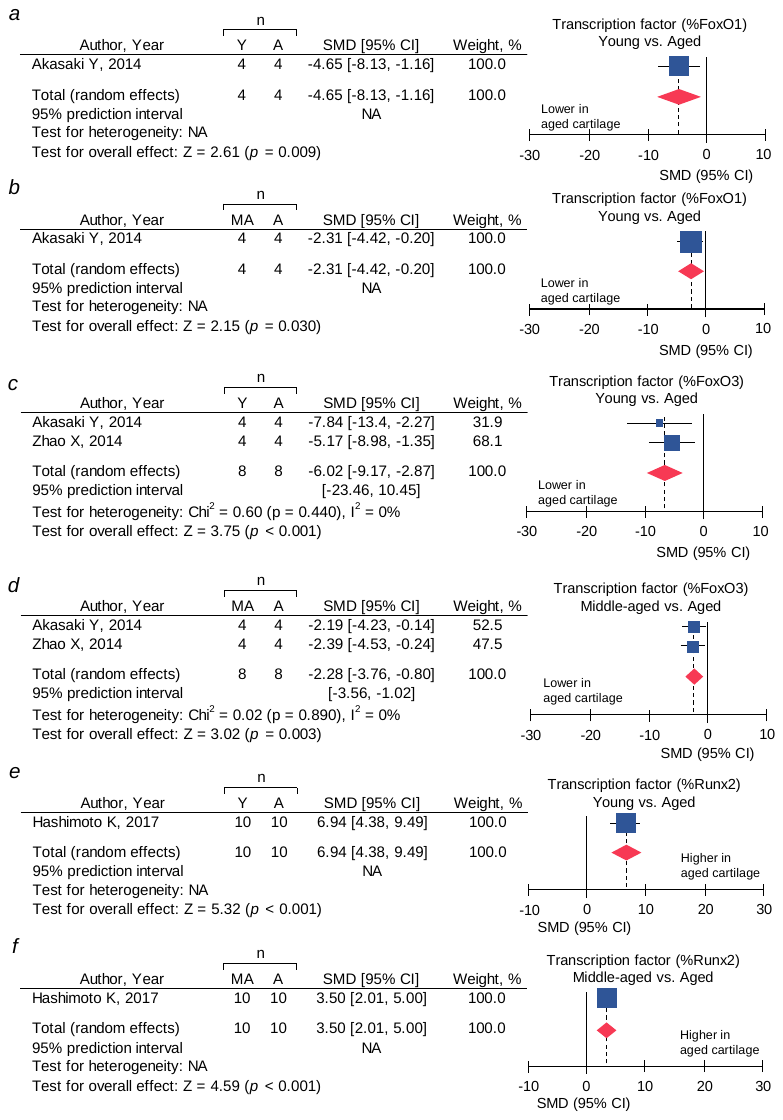

**Figure S8. SMD and 95% CI for transcription factor**

***a*.** %FoxO1-positive chondrocytes (young vs. aged). ***b*.** %FoxO1-positive chondrocytes (middle-aged vs. aged). ***c*.** %FoxO3-positive chondrocytes (young vs. aged). ***d*.** %FoxO3-positive chondrocytes (middle-aged vs. aged). ***e*.** %Runx2-positive chondrocytes (young vs. aged). ***f*.** %Runx2-positive chondrocytes (middle-aged vs. aged). The red diamond represents the pooled effect size (standardized mean difference; SMD). The vertical solid line at 0 represents no difference. The prediction interval is not displayed because of wide range. Y: young, MA: middle-aged, Y: aged.

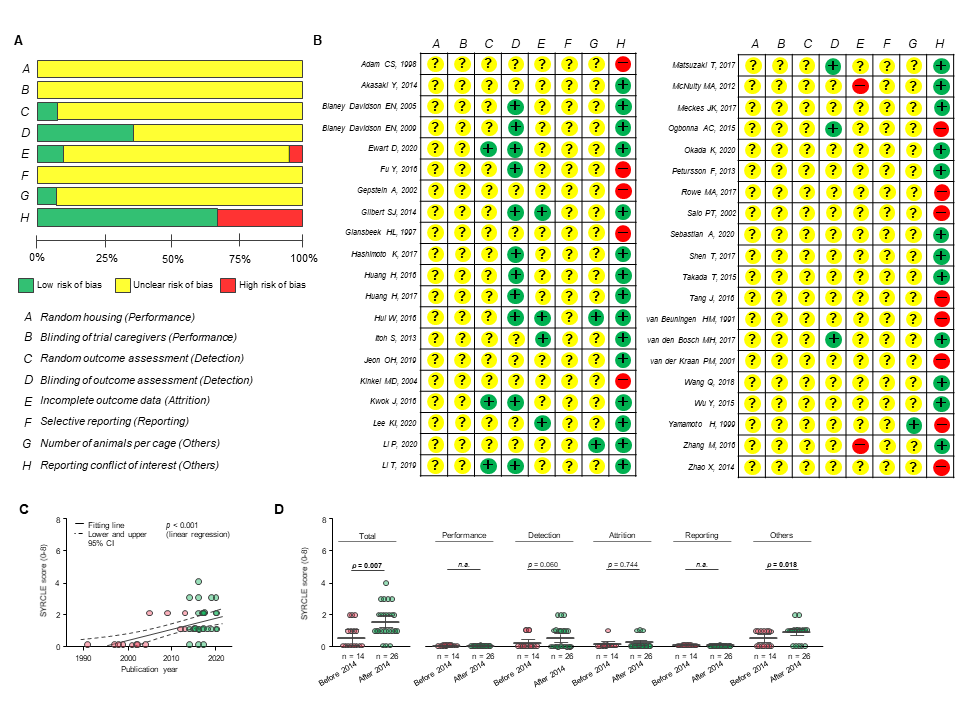

**Figure S9. Risk of bias assessment using SYRCLE**

**A.** Risk of bias of each item of SYstematic Review Centre for Laboratory animal Experimentation (SYRCLE). Each risk of bias item presented as percentages across all included studies, which indicates the proportion of different level risk of bias for each item. Publication that did not provide sufficient details to fulfill the criterion were judged as “unclear” in accordance with the original index of the SYRCLE(1). Overall, all domains considered using the SYRCLE, except for reporting conflict of interest, were judged as “unclear of risk of bias”. **B.** Risk of bias in individual studies. The red circle indicates high risk of bias, the green circle indicates low risk, and the yellow circle indicates that the risk is unclear as judged by 2 independent reviewers (HI and KW). Surprisingly, not one study adequately addressed item of random housing (performance bias), blinding of investigators (performance bias), or selective outcome reporting (reporting bias). Only 14 (35.0%) studies addressed blinding of outcome assessments (detection bias) that would be more important source of bias especially for subjective outcome assessment. **C.** Positive relationship between publication year and higher SYRCLE score (i.e., low risk of bias). Solid line represents fitting line and dotted lines represent lower and upper limits of 95% confidence interval. P-value of linear regression analysis is provided. **D.** Higher quality of reporting in studies published after publish of SYRCLE. Red (n = 14) and green dots (n = 26) represent independent studies published before and after publish of SYRCLE guideline (March 2014), respectively. Horizontal bars represent the means, and vertical bars represent the 95% confidence interval of the independent study. P-value was calculated using the two-sided Student t-test. We identified the evidence that the risk of bias is improving over time, which is attributed to improvement in detection bias (p = 0.060) and other bias (p = 0.018).

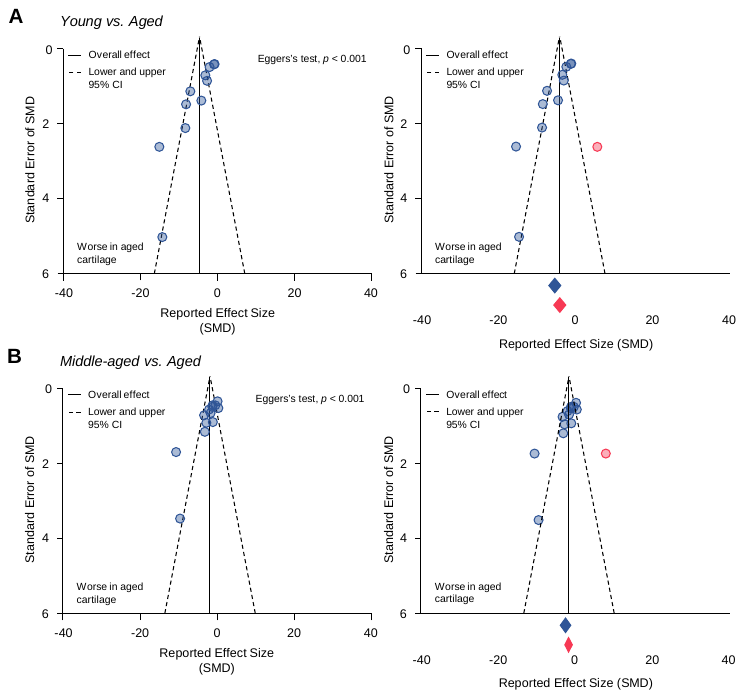

**Figure S10. Publication bias for cartilage morphology**

**A-B,** Funnel plot on age-related alteration in cartilage morphology compared to young (A) and middle-aged cartilage (B) before and after the trim-and-fill method. The vertical line represents the pooled standardized mean difference (SMD). The 2 diagonal dotted lines represent pseudo 95% confidence intervals around the summary effect for each standard error in the vertical axis. Red circle displays missing articles adjusted by the trim-and-fill method. The blue and red diamonds represent the observed and adjusted effect size, respectively. The degree of age-related cartilage degeneration was decreased (16.7% for vs. young, 26.0% for vs. middle-aged) after trim-and-fill-based adjustment for publication bias, which indicate that age-related cartilage degeneration is overestimated.

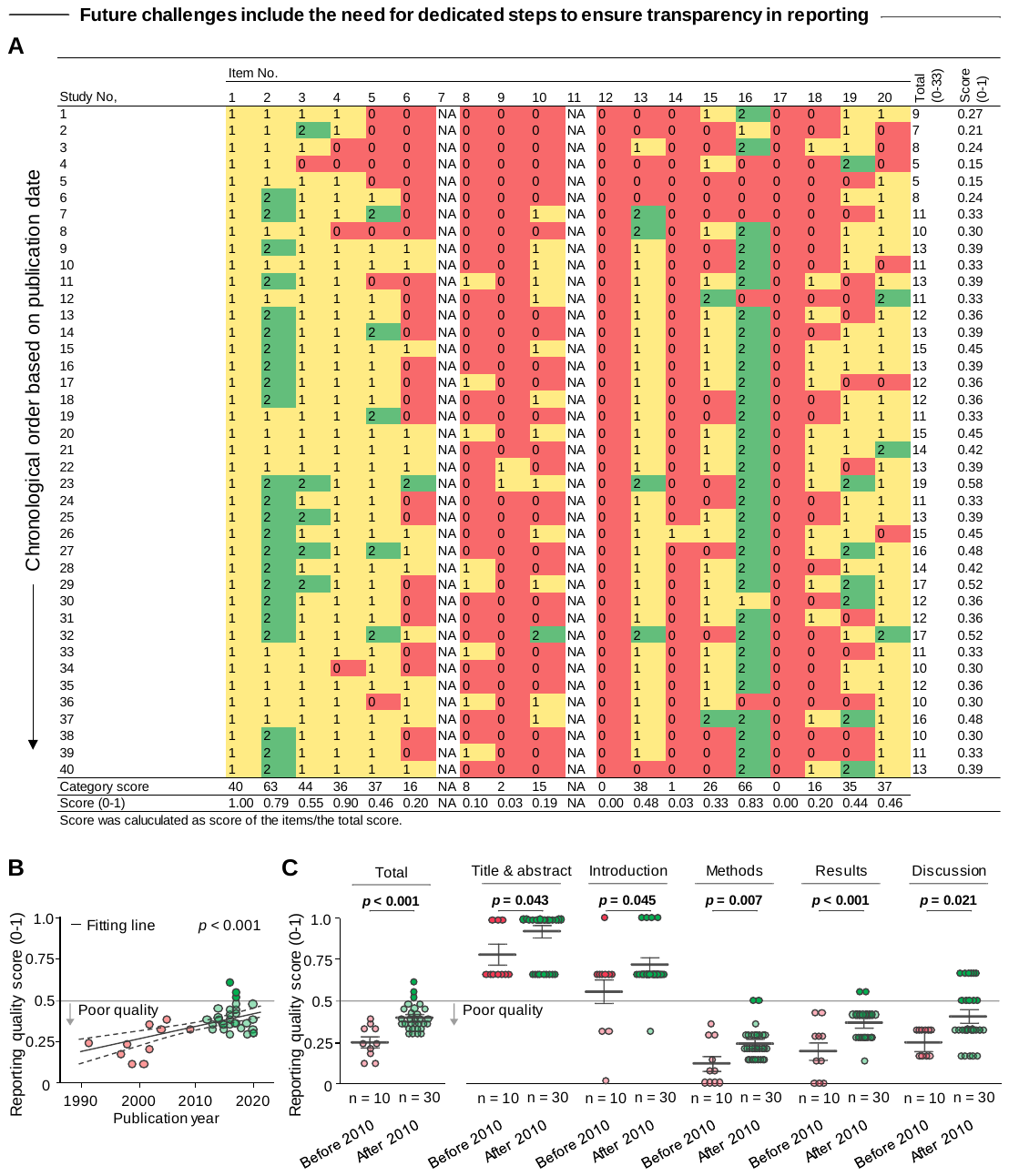

**Figure S11. Reporting quality of individual studies using ARRIVE guideline**

**A.** Heatmap of the ARRIVE guideline (reporting quality) of the included studies. Individual items indicate the following: (1) Title, (2) Abstract/Summary, (3) Introduction/Background, (4) Introduction/Primary and secondary objectives, (5) Methods/Ethical statement, (6) Methods/Study design, (7) Methods/Experimental procedure, (8) Methods/ Experimental Animals, (9) Methods/Housing and husbandry, (10) Methods/Sample size, (11) Methods/Allocation animals to experimental groups, (12) Methods/Experimental outcomes, (13) Methods/Statistical methods, (14) Results/Baseline data, (15) Results/Numbers analyzed, (16) Results/Outcomes and estimation, (17) Results/Adverse events, (18) Discussion/Interpretation and scientific implications, (19) Discussion/Generalizability and translation, and (20) Discussion/Funding. The heatmap was produced to facilitate the interpretation of results, where red indicates lowest score (0 point) of the category (clearly insufficient or unclear); yellow indicates middle score (1 point) of the category (sufficiency unclear); green indicates highest score (2 points) of the category (clearly sufficient or clear). **B.** Relationship between publication year and quality of reporting (reporting quality score, calculated as total score divided by maximum score [0-1]). Solid line represents fitting line and dotted lines represent lower and upper limits of 95% confidence interval. P-value of linear regression analysis is provided. **C.** Comparison of quality of reporting between studies published before (red, n = 10) and after (green, n = 30) publishing of ARRIVE (June 2010). Horizontal bars represent the means, and vertical bars represent the 95% confidence interval of the independent study. P-value was calculated using the two-sided Student t-test. Gray transverse line in (B) and (C) represents border (i.e., reporting quality score = 0.5) of the “poor” and “averaged” quality of evidence, and dark red and dark green dots represent studies that were judged as “averaged” quality of reporting.

**Supplementary methods**

This study was conducted according to the Preferred Reporting Items for Systematic reviews and Meta-Analyses (PRISMA) statement(3) (**Supplemental Appendix S3**), PRISMA protocols (PRISMA-P)(4), Meta-analysis of Observational Studies in Epidemiology (MOOSE) checklist(5) (**Supplemental Appendix S4**), Cochrane Handbook for Systematic Reviews of Interventions(6), and the practical guide for meta-analysis from animal studies(7).

***Eligibility criteria***

Manuscript eligibility criteria were defined according to the PECO. In brief, we included articles characterizing aged articular cartilage in the knee joints of mice (i.e., each study had to include both aged and young mice or aged and middle-aged mice). Young, middle-aged, and aged mice were defined as 2-7, 9-15, and 18-24 months old, respectively, and correspond to ages 20-30, 38-47, and 56-69 years in humans, respectively(8).

This review was limited to mouse models, as this is the preferred model given their relatively fast disease progression, cost-efficiency, and ease of handling(9). Rats, on the other hand, are generally free of spontaneous OA(10). Furthermore, several assessment tools, including micro-computed tomography (CT), gait analysis, and pain assessment are now available for mouse models, thereby allowing for a more comprehensive assessment of pathology(11). No restrictions were set according to mouse strain, or manuscript publication year. The following articles were excluded: (1) studies that included genetically modified animals, as such models likely oversimplify the disease process, whereas naturally occurring OA is almost certainly polygenic in nature(12); and (2) studies that did not explicitly compare knee articular cartilage between aged and young mice or aged and middle-aged mice.

Outcome measurements related to assessment of articular cartilage damage were divided into seven principal outcome categories based on a slightly modified version of previous meta-analyses(13, 14). Outcome measures in pre-clinical studies should be directly relevant to the human disease/intervention(15) and should be minimally invasive(16). This review included age-related changes of molecular biomarker in blood, urine, and synovial fluids as an outcome variable, as these are promising disease markers in predicting structural OA progression(17). We used the following hierarchy of articular cartilage-related outcomes:

(1) Morphology (e.g., Osteoarthritis Research Society International [OARSI] score(18), Mankin’s score(19), MRI based classifications of morphological changes);
(2) morphometry (any kind of quantitative methods on microscopic images of articular cartilage including computational image analysis techniques(20));
(3) Composition (any kind of quantitative methods for quantification of proteoglycan or collagen);
(4) biomechanical characterization (e.g., tensile and compressive measures of stiffness);
(5) biomarker (e.g., mirco RNA in synovial fluid);
(6) molecular biology (e.g., ECM-related gene expression and protein synthesis); and
(7) others (e.g., cartilage composition, pain behavior, gait analysis(21)).

***Literature search***

PubMed, Physiotherapy Evidence Database (PEDro), Cumulative Index to Nursing and Allied Health Literature (CINAHL), and Cochrane Central Register of Controlled Trials electronic databases were used. Google Scholar was also used as a complementary search engine. A manual search of the reference lists of past reviews was performed(9, 11, 16, 22-32). Furthermore, citation searching was performed on the original record using Web of Science. These citation indexes are recommended by the Cochrane Handbook(6). For each electronic database, the database search was performed on March 10, 2020. Electronic searches used combined key terms, including “Animal, Laboratory” “Aging,” “Age factors,” “Cartilage, Articular,” and “Osteoarthritis, Knee” using Medical Subject Headings terms.

***Study selection***

Two independent reviewers (HI and GG) assessed eligibility in accordance with the Cochran Handbook recommendations(6). Two reviewers screened titles and abstracts yielded by the search. Full manuscripts of the articles that met the eligibility criteria were then obtained and reviewed. During these processes, the reviewers prepared and used simple, pre-designed Google spreadsheets to assess eligibility by extracting study features. Disagreements between the two reviewers were discussed until consensus was achieved.

***Data collection***

A single reviewer (HI) extracted data regarding basic study information (authors, publication year, and country of corresponding author), experimental condition (i.e., mice strain, age, sample size, and sex), target joint (tibiofemoral or patellofemoral joints), outcome measures, funding, and presence of conflict of interest. These data were extracted because of their potential to influence key outcome measures (except for basic study information)(33, 34). If outcome measures from multiple time points were reported within same age category (e.g., three and six months old from the young group), we averaged the effect size (35). If outcome measures from multiple compartments (e.g. medial and lateral compartments in tibiofemoral joint and patellofemoral joint) were presented, data from the most severe region was extracted. When data were not reported or unclear, we contacted the authors directly. A reminder was sent to those who had not replied. If data were provided only in figures, the graphically presented data was converted to numerical data using a reliable and validated digital ruler software (WebPlotDigitizer)(36, 37). To further confirm the reliability of the data converting, another independent reviewer (SS) re-calculated the data from 5 randomly selected articles. Inter-rater reliability between two reviewers was excellent (intraclass correlation coefficient [2,1]: 0.999)(38).

***Synthesis of results***

*Meta-analysis*

To characterize aged articular cartilage and the underlying mechanism of age-related cartilage degeneration, pooled estimates and 95% confidence intervals for standardized mean differences (SMD) of outcome measures were calculated using the DerSimonian-Laird method(39). This method considers the precision of individual studies and the variation between studies and weighs each study accordingly. SMD were calculated using the mean between-group difference (aged and middle-aged or young) divided by the pooled standard deviation(39). Meta-analyses were performed using Review Manager Version 5.3 (Nordic Cochrane Centre, Cochrane Collaboration, Copenhagen, Denmark). Prediction intervals for each outcome variable were also estimated(40). In the synthesis, mean or SD values of 17 studies(41-57) were converted from graphical data by reliable software as described above. In one study(45), sample size in the young, middle-aged, and aged groups were provided by the primary author. Study heterogeneity, defined as the inter-trial variation in study outcomes, was assessed using *I^2^*, which is the proportion of total variance explained by inter-trial heterogeneity(58).

To address the trajectory of age-related changes in articular cartilage, a mixed linear regression analysis with random slopes and random intercepts was performed for the outcome of cartilage morphology using SPSS Statistics for Windows, version 25.0 (IBM Corp., Armonk, NY, USA). In this analysis, age category (1: young, 2: middle-aged, 3: aged) and standardized semi-quantitative score of cartilage degeneration was included as independent and dependent variables, respectively. To standardize semi-quantitative score of cartilage degeneration, all histological scores provided in each included study were converted to 0-100 and recalculated as in a previous meta-analysis(59), with higher score indicates severe cartilage degeneration.

*Functional characterization of transcriptome*

Single sample gene set enrichment analysis (ssGSEA) was performed by R/Bioconductor package fgsea with gene scores defined by log2 fold change of gene expression profiles among groups of young, middle-aged, and aged that is available from one study(60) included in this systematic review. The same analyses were performed using transcripts from PTOA model that is available from the same study(60). We used GO terms (GO_Biological_Process_2018) as a gene set downloaded from enrichr (https://amp.pharm.mssm.edu/Enrichr/). REVIGO software(61) was applied to summarized redundant GO terms and visualize the summarized results. Transcription factor enrichment analysis was also performed using ChIP-X Enrichment Analysis Version 3 (ChEA3)(62) from the RNA-seq data with false discovery rate adjusted p-value less than 0.05.

*PPI network construction and pathway enrichment analysis*

The Search Tool for the Retrieval of Interacting Genes database (STRINGdb)(63) was used for the construction of the PPI network based on differentially expressed (p<0.05) proteins. The type of connection included the following items: textmining, experiments, databases, co-expression, neighborhood, gene fusion, and co-occurrence. To extract valid interactions, a confidence score of >0.4 was set as the cut-off criterion. Kyoto Encyclopedia of Genes and Genomes (KEGG) pathway enrichment analyses were conducted using the STRINGdb(63).

***Overall quality of evidence***

Two independent reviewers (HI and KW) performed assessment of reporting quality (ARRIVE)(64), risk of bias (SYRCLE)(1), publication bias, and quality of evidence (GRADE)(2) in a blinded manner. Disagreements between two reviewers were discussed until consensus was achieved.

*Reporting quality: ARRIVE*

The reporting quality of all included *in vivo* studies in this systematic review was assessed according to the modified version of ARRIVE (Animal Research: Reporting In Vivo Experiments) guidelines. The reporting quality was evaluated on the basis of a predefined ARRIVE grading system(65) applied to the following items: (1) Title, (2) Abstract/Summary, (3) Introduction/Background, (4) Introduction/Primary and secondary objectives, (5) Methods/Ethical statement, (6) Methods/Study design, (7) Methods/Experimental procedure, (8) Methods/ Experimental Animals, (9) Methods/Housing and husbandry, (10) Methods/Sample size, (11) Methods/Allocation animals to experimental groups, (12) Methods/Experimental outcomes, (13) Methods/Statistical methods, (14) Results/Baseline data, (15) Results/Numbers analyzed, (16) Results/Outcomes and estimation, (17) Results/Adverse events, (18) Discussion/Interpretation and scientific implications, (19) Discussion/Generalizability and translation, and (20) Discussion/Funding. The modified version does not include items (7) and (11), as these items involve interventional studies and are therefore not applicable. Furthermore, the two reviewers did not consider strain of animal used in the abstract (item 2), as addition of this information in the abstract is not realistic for those studies that used several mouse strains within a strict word limit. The maximum score by column was calculated to obtain the quality score. For descriptive purposes, we calculated the standardized score (quality score/maximum score) by generating three possible range coefficients in which 0.8–1 was considered “excellent”, 0.5–0.8 was considered “average”, and scores below 0.5 were considered “poor”(66). To evaluate the relationship between the publication year and standardized ARRIVE score, linear regression analysis was performed. We checked the normal distribution of the data using Shapiro-Wilk test, and the features of the regression model by comparing the residuals vs. fitted values (i.e., the residuals had to be normally distributed around zero). To probe whether publication of the ARRIVE guidelines affected reporting quality, a post-hoc two-sided Student t-test was used to compare the standardized score before and after publication of the ARRIVE guideline (June, 2010). Although the ARRIVE guidelines were recently updated in 2020, we chose to use the 2010 guidelines so that we could adequately group studies based on before and after their publication.

*Risk of bias: SYRCLE*

This study used the modified version of SYstematic Review Centre for Laboratory animal Experimentation’s (SYRCLE’s) tool(1) to assess the risk of bias in each study. This assessment tool is based on the Cochrane risk of bias tool(67) and has been adjusted for animal studies. We did not apply items of random sequence generation, baseline characteristics, and allocation concealment, as these items are not applicable to the included studies. The modified version of the SYRCLE consists of five types of bias (performance, detection, attrition, reporting, and other) across the following eight domains: (1) random housing; (2) blinding (caregivers and/or investigators); (3) random outcome assessment; (4) blinding (outcome assessor); (5) incomplete outcome data (i.e., missing outcome data); (6) selective outcome reporting; (7) number of animals per cage; and (8) statement of potential conflict of interest. Judgements for (7)-(8) were made because these factors may influence the outcome measures(33, 68-70). The score “yes (+)” indicates a low risk of bias, “no (-)” indicates high risk of bias, and “?” indicates an unclear risk of bias. We used the signaling questions provided in the original article wherever possible(1). Concerning incomplete outcomes (attrition bias; domain 5), we assumed that no exclusions were made if the number of animals per group mentioned in the materials and methods section was identical to the number stated in the figure legends or results section. To evaluate the relationship between the publication year and standardized SYRCLE score, linear regression analysis was performed. We checked the normal distribution of the data using Shapiro-Wilk test, and the features of the regression model by comparing the residuals vs. fitted values (i.e., the residuals had to be normally distributed around zero). To probe whether publication of the SYRCLE affected risk of bias, a post-hoc two-sided Student t-test was used to compare the standardized score before and after publication of the SYRCLE guideline (March 2014).

*Publication bias*

We evaluated publication bias for outcome measures only if more than 10 studies were synthesized into meta-analysis(71). We used a linear regression approach, Egger’s regression, to measure funnel plot asymmetry (i.e., publication bias). The Egger’s test for funnel plot asymmetry formally examines whether the association between estimated effects size and a measure of standard error of the effects size is greater than might be expected to occur by chance(72). A value of *p* < 0.100 indicated the existence of publication bias.

*Quality of evidence: GRADE*

The Grading of Recommendations Assessment, Development, and Evaluation (GRADE) approach was used to assess the quality of evidence for each outcome measure(2, 73). GRADEpro GDT (GRADEpro Guideline Development Tool [Software]. McMaster University, 2015 (developed by Evidence Prime, Inc.)) was used to import data from the Review Manager to summarize findings in a table. The methodological criteria by which evidence was assessed depended on five primary domains (risk of bias, inconsistency, indirectness, precision, and publication bias). The evidence quality was downgraded if (1) outcomes have a high risk of bias; we defined this as a lack of blinding outcome assessment in more than 50% of the included studies (risk of bias domain); (2) heterogeneity between trials was more than substantial (*I^2^* ≥ 50%) with non-overlapping 95% CI(6, 74) (inconsistency domain); (3) sample size was inadequate; we defined this as optimal information size(75) and wide 95% CI that included 0 (precision domain); and (4) publication bias existed, as identified by the Egger’s regression test (publication bias domain). The publication bias domain was applied only for outcome with ≥10 studies. Downgrade in the indirectness domain was not applied, as all studies used mouse models and outcomes were not directly related to human clinical trials and clinical decisions. The quality of evidence was judged as “high”, “moderate”, “low,” or “very low”.
