## Supplemental Appendix S4 for "Meta-analysis and multi-omics to elucidate pathogenic mechanisms of age-related knee osteoarthritis"

A Proposed Reporting Checklist for Authors, Editors, and Reviewers of Meta-analyses of Observational Studies.

| **#** | **Checklist item** | **Reported on page #** |
| --- | --- | --- |
| **Reporting of background should include** | |  |
| 1 | Problem definition | 4-5 |
| 2 | Hypothesis statement | Not applicable |
| 3 | Description of study outcome(s) | 4-5 |
| 4 | Type of exposure or intervention used | 4-5 |
| 5 | Type of study designs used | 5 |
| 6 | Study population | 4-5 |
| **Reporting of search strategy should include** | |  |
| 7 | Qualifications of searchers (eg., librarians and investigators) | Suppl. methods |
| 8 | Search strategy, including time period included in the synthesis and keywords | 16-17, Suppl. methods |
| 9 | Effort to include all available studies, including contact with authors | Suppl. methods |
| 10 | Databases and registries searched | Suppl. methods |
| 11 | Search software used, name and version, including special features used (eg., explosion) | Suppl. methods |
| 12 | List of citations located and those excluded, including justification | Not applicable |
| 13 | Method of addressing articles published in languages other than English | Not applicable |
| 14 | Method of handling abstracts and unpublished studies | 16, Suppl. methods |
| 15 | Description of any contact with authors | Suppl. methods |
| **Reporting of methods should include** | |  |
| 16 | Description of relevance or appropriateness of studies assembled for assessing the hypothesis to be tested | Not applicable |
| 17 | Rationale for the selection and coding of data (eg., sound clinical principles or convenience) | Suppl. methods |
| 18 | Documentation of how data were classified and coded (eg., multiple raters, blinding, and interrater reliability) | Suppl. methods |
| 19 | Assessment of confounding (eg., comparability of cases and controls in studies where appropriate) | Not applicable |
| 20 | Assessment of study quality, including blinding of quality assessors; stratification or regression on possible predictors of study results | 17  Suppl. methods |
| **#** | **Checklist item** | **Reported on page #** |
| 21 | Assessment of heterogeneity | 17, Suppl. methods |
| 22 | Description of statistical methods (eg., complete description of fixed or random effects model, justification of whether the chosen models account for predictors of study results, dose-response models, or cumulative meta-analysis) in sufficient detail to be replicated | 17, Suppl. methods |
| 23 | Provision of appropriate tables and graphics | Suppl. methods |
| **Reporting of results should include** | |  |
| 24 | Graphic summarizing individual study estimates and overall estimate | Fig.2, Suppl. Figure |
| 25 | Table giving descriptive information for each study included | Table 1 |
| 26 | Results of sensitivity testing (eg., subgroup analysis) | Not applicable |
| 27 | Indication of statistical uncertainly of findings | 11, Table 2 |
| **Reporting of discussion should include** | |  |
| 28 | Quantitative assessment of bias (eg., publication bias) | 12, Suppl. Figure |
| 29 | Justification for exclusion (eg., exclusion of non-English-language citations) | Suppl. Table |
| 30 | Assessment of quality of included studies | 12, Suppl. Figure |
| **Reporting of conclusions should include** | |  |
| 31 | Consideration of alternative explanations for observed results | 13-16 |
| 32 | Generalization of the conclusions (ie., appropriate for the data presented and within the domain of the literature review) | 13-16 |
| 33 | Guidelines for future research | 13-16 |
| 34 | Disclosure of funding source | 17 |

From: Stroup DF, Berlin JA, Morton SC, et al. Meta-analysis of observational studies in epidemiology: a proposal for reporting. Meta-analysis Of Observational Studies in Epidemiology (MOOSE) group.

JAMA 2000;283(15):2008–12.
